# A Genome-Phenome Association study in native microbiomes identifies a mechanism for cytosine modification in DNA and RNA

**DOI:** 10.1101/2021.03.23.436658

**Authors:** Weiwei Yang, Yu-Cheng Lin, William Johnson, Nan Dai, Romualdas Vaisvila, Peter R. Weigele, Yan-Jiun Lee, Ivan R. Corrêa, Ira Schildkraut, Laurence Ettwiller

## Abstract

Shotgun metagenomic sequencing is a powerful approach to study microbiomes in an unbiased manner and of increasing relevance for identifying novel enzymatic functions. However, the potential of metagenomics to relate from microbiome composition to function has thus far been underutilized. Here, we introduce the Metagenomics Genome-Phenome Association (MetaGPA) study framework, which allows to link genetic information in metagenomes with a dedicated functional phenotype. We applied MetaGPA to identify enzymes associated with cytosine modifications in environmental samples. From the 2365 genes that met our significance criteria, we confirm known pathways for cytosine modifications and proposed novel cytosine-modifying mechanisms. Specifically, we characterized and identified a novel nucleic acid modifying enzyme, 5-hydroxymethylcytosine carbamoyltransferase, that catalyzes the formation of a previously unknown cytosine modification, 5-carbamoyloxymethylcytosine, in DNA and RNA. Our work introduces MetaGPA as a novel and versatile tool for advancing functional metagenomics.

## Introduction

Advances in next-generation sequencing technology have been reshaping metagenomics, making it possible to explore all microbes within a sample, including the overwhelming majority of those unculturable (Quince et al., 2017). From studying human gut microbiome to marine viral communities (Sunagawa et al., 2015), metagenomic analyses are being utilized across a broad spectrum of life science disciplines, contributing to novel clinical diagnoses (Chiu and Miller, 2019), antibiotics and small molecule discovery (Charlop-Powers et al., 2014; Hu et al., 2013), food safety (Cao et al., 2017), biofuel generation (Hess et al., 2011) and environment stewardship (Cavicchioli et al., 2019; Nesme et al., 2014).

Of the two primary questions metagenomic studies strive to address, profiling the taxonomic biodiversity – “who is out there?” is easier to answer compared to inferring the biological functions – “what do they do?”. Indeed, there have been many well-established strategies to quantify taxonomic diversity such as analyzing known marker genes, binning contigs and assembling sequences into taxonomic groups or genomes (Quince et al., 2017; Sharpton, 2014). However, a large fraction of the functional potential of microbiomes remains to be discovered. Microbiomes encode a large number of evolutionarily diverse genes coding for millions of peptides and proteins for distinctive functions. The functional diversity is particularly enormous for secondary metabolites and epigenetic modifications (Sharon et al., 2014).

Taking DNA modifications as an example of functional diversity, environmental microbiomes are a plentiful source of DNA-modifying enzymes with diverse mechanisms (Weigele and Raleigh, 2016) (Hiraoka et al., 2019). In bacteriophages, DNA modifications are used to evade the restriction-modification systems of the host bacterial cell. Notably, a handful of bacteriophages have been described to completely modify cytosines in their genomic DNA, e.g. 5’ methylcytosine in XP12 phage (Kuo et al., 1968) and glycosylated 5’ hydroxymethylcytosine in T4 phage (Revel and Georgopoulos, 1969). Recently, additional base modifications have been discovered including 5-(2-aminoethoxy)methyluridine, 5-(2-aminoethyl)uridine and 7-deazaguanine (Hutinet et al., 2019; Lee et al., 2018).

Current methods to harness such functional potential of microbiomes, for the most part, came from computational prediction of gene function based on homology search to existing databases. Nonetheless, because of the poor completeness and accuracy of microbial annotation, homology searches often fail to impute the correct functions (Huson et al., 2009; Nayfach and Pollard, 2016). Novel functions can also be found using functional screens; however, such effort entails the construction and high-throughput screening of a large number of clones which can be very time-consuming (Martinez et al., 2004). Novel approaches are therefore needed to link genetic information from microbial metagenomes to function.

Here we developed the Metagenomics Genome-Phenome Association (MetaGPA) framework to bridge the gap between genetic information and functional phenotype. MetaGPA is conceptually close to Genome-Wide Association Studies (GWAS) as it associates genotypic data with phenotypic traits. We incorporated association analyses at the level of protein domains to reveal genes significantly associated with the phenotype of interest. By applying this workflow on DNA modifications as our phenotypic trait, we discovered a number of candidate enzyme families. From these candidates, we validated a novel DNA/RNA cytosine modification, 5-carbamoyloxymethylcytosine, and the enzyme responsible for this modification. From this example, we show that MetaGPA is a powerful and versatile method to improve metagenome functional analysis.

## Result

### 1. Conceptual framework of Metagenomics Genome-Phenome Association (MetaGPA) studies

Microbiomes are, for a large part, unculturable and studies in microbiome composition are strongly influenced by laboratory culture conditions. It is therefore challenging to study their native functions through classical genetics. We hypothesize that MetaGPA provides an alternative by associating genetic units within microbiomes at genome-scale with distinct phenotypic readouts (**Fig. 1**).

**Figure 1:**
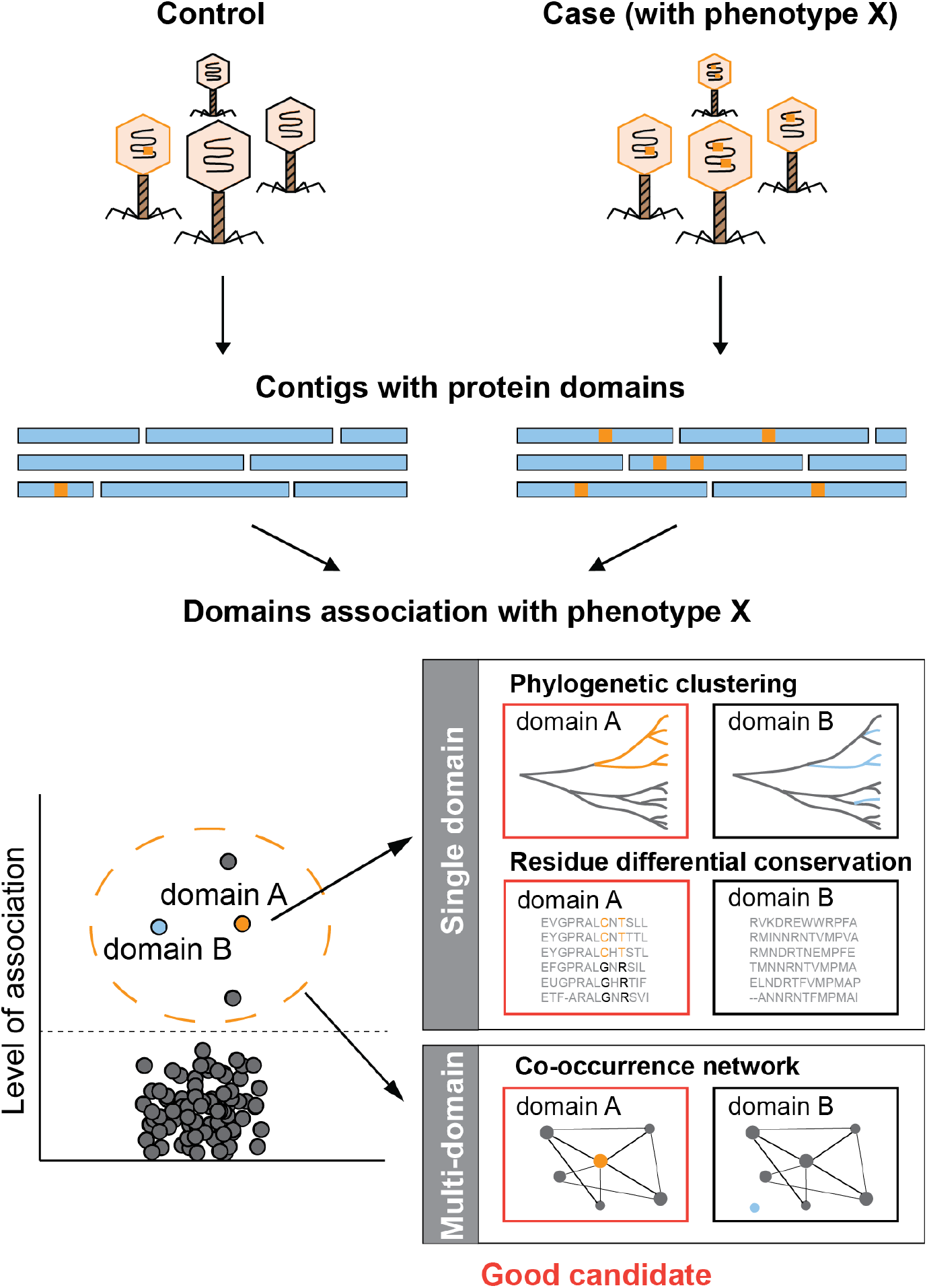
Schematic overview of MetaGPA. MetaGPA is applied to identify protein domain families that are significantly associated with a given phenotype (in orange bar). It performs association analyses at single and multi-unit levels to highlight functional units that are most likely associated with the selected phenotype. In this theoretical example, colored branches or residues (orange or light blue) represent contigs from the case cohort. Both domain A and B are associated with the given phenotype but the phylogenetic clustering and co-occurrence network of their candidate genes prioritize domain A. More detail about the workflow is provided in **Supplementary Fig. 1**.

Like GWAS in individual species (Hirschhorn and Daly, 2005), MetaGPA requires definition of two cohorts, a “case” cohort, i.e., a group of organisms that share a specific phenotype under study, and a “control” cohort composed of all organisms within the same biological specimen (**Fig. 1** and **Supplementary Fig. 1**). Both cohorts are sequenced, *de-novo* assembled into contigs and genes identified in these contigs are annotated using homology search to known protein domains. Genes in the case cohort containing protein domain families that are significantly associated with the phenotype define candidate genes.

Both protein domain families and their corresponding candidate genes can be further refined using evidence such as phylogenetic clustering and co-occurrence with other candidate domain families/genes. Phylogenetic clustering assesses for each associated domain, whether candidate genes are phylogenetically closer to each other relative to genes containing the same domain family in the control cohort. This analysis can be done at domain or even residue resolution. Co-occurrence with other candidates genes strengthen the association by attempting to identify the entire metabolic pathway responsible for the specific phenotype under study. Taken together, this multilayer analysis effectively identifies protein domain families and their corresponding candidate genes that are truly related to the phenotypes of interest.

### 2. Application of MetaGPA for the discovery of cytosine modifying and hyper-modifying enzymes

#### Principle

To demonstrate the effectiveness of the MetaGPA framework, we designed a study to identify novel proteins associated with DNA cytosine modifications. Because the relevant phenotype (cytosine modifications) is covalently linked to the genetic material, total genomic DNA isolated from environmental sources can be used for both, the ‘control’ cohort and material for phenotypic selection.

Accordingly, a ‘case’ cohort is obtained by applying an enzymatic selection to retain only the genomic DNA containing known and unknown forms of cytosine modification. More specifically, unmodified cytosines are deaminated to uracils using the DNA cytidine deaminase Apolipoprotein B mRNA editing enzyme catalytic polypeptide-like 3A (APOBEC3A) (Carpenter et al., 2012), and subsequently excised by Uracil-Specific Excision Reagent (USER) (Bitinaite et al., 2007), resulting in fragmented DNA. Prior to APOBEC3A treatment, ten-eleven translocation dioxygenase 2 (TET2) and T4 phage β-glucosyltransferase (T4-BGT) can be used to protect 5-methyl-2’-deoxycytidine (5mdC) and 5-hydroxymethyl-2’-deoxycytidine (5hmdC) from deamination and later excision (**Fig. 2a**).

**Figure 2:**
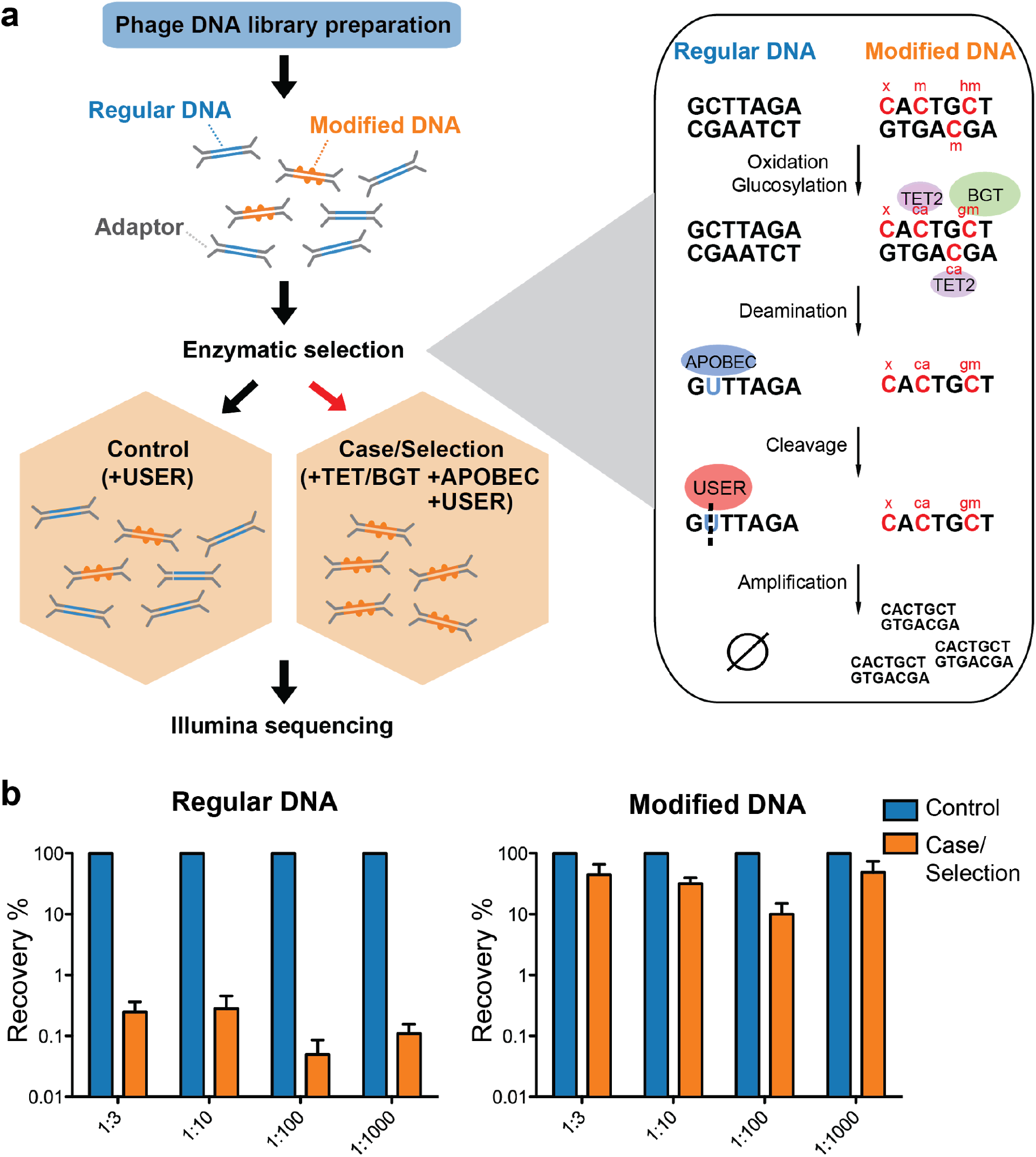
Selective sequencing of DNA containing modified cytosines. **a**, Schematic of enzymatic selection. Adapter ligated products were first incubated with TET2 and T4-BGT so that 5mdC (m) and 5hmdC (hm) in these sequences may be oxidized to 5-carboxycytosine (ca) or glucosylated to 5-β-glucosyloxymethylcytosine (gm) (**Materials and Methods**). Unmodified cytosines were deaminated by APOBEC3A into uracils and then cut by USER, so that DNA with modified cytosines is enriched after the selection. This method is predicted to also preserve unknown forms of cytosine modification (denoted “x”) provided that they block C-to-U deamination. **b**, Enrichment of modified DNA with serial dilutions. A total of 250 ng genomic DNA (from *E. coli)* and cytosine-modified genomic DNA (from T4gt) were used for enzymatic selection or control per assay. Bar graphs show average recovery of DNA from three independent experiments ± SEM.

Effectively, unmodified DNA is depleted and the remaining material in the ‘case’ cohort should only comprise DNA containing modified cytosines resistant to APOBEC3A treatment. We hypothesize that many forms of cytosine modification, including those unknown to date, are naturally protected from deamination by APOBEC3A, and should be amenable to enrichment by our selection method.

#### Cytosine modified DNA is retained after enzymatic selection

To demonstrate the feasibility of this approach, a mock community consisting of an equimolar amount of genomic DNA from *E. coli* (containing unmodified cytosine, dC) and T4gT phage (containing 90-95% 5hmdC) was sheared, subjected to enzymatic selection and qPCR was performed to quantify the remaining DNA (see **Materials and Methods**). Compared to the original mock community, enzymatic selection resulted in a 0.5% recovery of *E. coli* DNA and an average 35% recovery of T4gT DNA (**Fig. 2b**). This result shows that DNA containing modified cytosine is about seventy-fold enriched compared to unmodified DNA.

To test the sensitivity and efficiency of this method, we serially diluted modified DNA and unmodified DNA to 1:3, 1:10, 1:100 and 1:1000 molar ratio and carried out the enzymatic selection. Recovery rates were calculated and compared to no-enzyme treatment control. Even at 1:1000 dilution, an average of 48.6 % modified DNA was retained relative to no-enzyme control. This result showed the capability of our method to concentrate trace amounts (picogram-level) of modified DNA from a complex sample (**Fig. 2b**, **Supplementary table 1**).

#### MetaGPA association analyses on complex microbiomes identifies candidate enzymes with DNA modification potential

We collected environmental samples for the aim of identifying novel enzymes involved in cytosine modification using MetaGPA. To explore the robustness of MetGPA, we included 3 environmental samples from distinct sources (**Fig. 3a**); of the three samples, two were collected at a sewage treatment plant and the other one was collected from coastal sludge.

**Figure 3:**
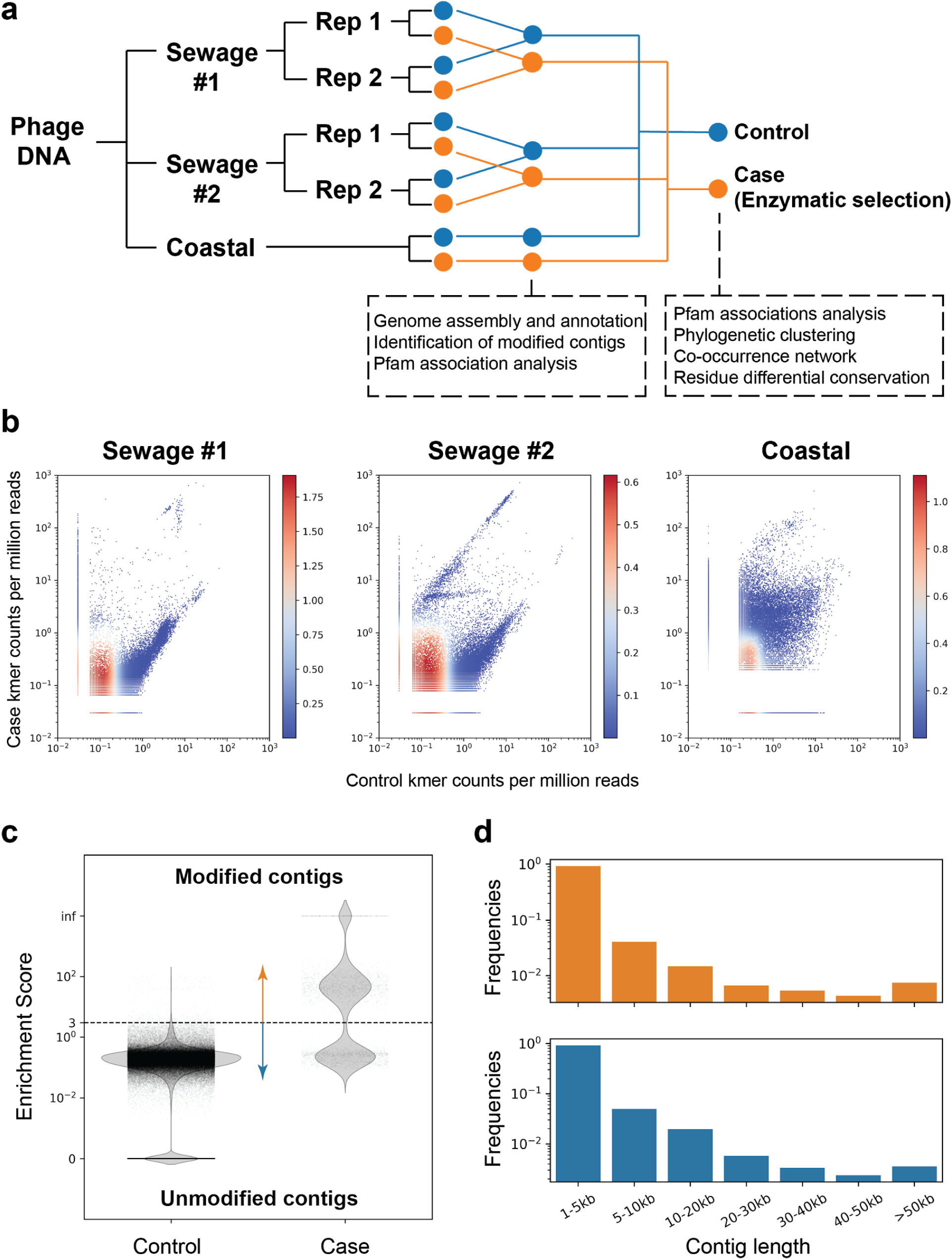
Screening of contigs associated with DNA modification. **a**, Overview of the environmental samples and analysis used in this study. Three independent sequencing experiments were performed on three samples, two sewage samples (same sampling location at different times #1 and #2) and one coastal sample (Beverly, MA). Phage DNA was collected and extracted and for the sewage samples, two technical replicates (Rep 1 and 2) were performed. Analyses performed with separate or composite datasets were indicated in below blocks. **b,** Plots of k-mer composition. Plots show normalized 16-mer count density with randomly selected subsamples of 0.1% of the data from each experiment. Red in the colorbar represents high density while blue represents low density. **c**, Dotplot represents the enrichment scores for each non-redundant contig in the control (left) and case/enzymatic selection (right) cohorts. Dashed line separates modified from unmodified contigs based on a threshold enrichment score equaled to 3 (Enrichment score = RPKM_(selection)_ / RPKM_(control)_). The plot shows every contig from all three environmental samples. **d,** Distributions of modified (orange) and unmodified (blue) contigs length (in kilobases, kb).

We aim at obtaining the phage-enriched fraction as phages are known to carry a large diversity of modifications (Weigele and Raleigh, 2016). For this, each sample was precipitated with polyethylene glycol (PEG) followed by DNA extraction using phenol/chloroform (see **Materials and Methods**). Libraries were made in duplicates (except for the coastal sample) using modified adapters to protect them from enzymatic degradation. Libraries were subjected to either enzymatic selection (i.e. case) or no enzymatic selection (i.e. control) prior to amplification (**Fig. 3a**). Additionally, spiked-in genomic DNA mixture of *E. coli* (dC), XP12 (5mdC) and T4gt (5hmdC) were added to each sample after adapter ligation and before enzymatic treatment. Recovery of spiked-in modified DNAs was detected as expected (**Supplementary Fig. 2a**). Accordingly, we proceeded with Illumina high throughput sequencing and obtained an average of 60 million paired-end reads per library. Analysis of the read composition reveals consistency of k-mer composition between replicates, demonstrating that our enzymatic selection for modified DNA is reproducible (**Supplementary Fig. 2b**). Reads from both replicates were combined and normalized k-mer frequency plots showed diversity of k-mer composition from different sources/samples, while highlighting a small portion of k-mers that was highly enriched in the selection libraries (**Fig. 3b**). To translate and study the biological entities from the dataset, we separately assembled the reads from the case (selection) and control datasets from the three environmental samples into contigs and removed contigs that were either too short (less than 1000 bp) or redundant (**Materials and Methods**). Reads were mapped to the contigs and the ratio between the normalized coverage in the case (selection) library (RPKM_(case)_) and the normalized coverage in the control library (RPKM_(control)_) defines the enrichment score for each contig (**Materials and Methods**, **Supplementary Fig. 2c**). A high enrichment score suggests that the contig is derived from DNA containing modified cytosine (i.e. modified contig). We define as modified, contigs with an enrichment score equal or above 3. In total, 3901 modified contigs were identified from three DNA samples (**Fig. 3c** and **Supplementary Fig. 2c-d**). Distribution of contigs across all length ranges was equal between modified and unmodified contigs (**Fig. 3d**) and the proportion of modified contigs among sewage and coastal samples were comparable (about 2% in both sewage samples and 3.5% in coastal total contigs) (**Supplementary Fig. 2d**).

Annotations of protein domain families were performed using profile hidden Markov models of all protein domain families described in the Pfam database(El-Gebali et al., 2019)(**Materials and Methods**). Instances of each domain family in the modified and unmodified contigs were calculated and Fisher’s exact test was conducted to reveal domain families that were positively associated with DNA modification. Interestingly, there was a high degree of overlap between top associated domain families among different environmental samples (**Supplementary Fig. 3a**), suggesting that a group of universal protein families for DNA modification may exist regardless of the origin of the microbiome. Given this consistency, we decided to pool the reads from three environmental samples together and repeat the association analysis to achieve higher statistical power. Associated domains were subsequently matched to open reading frames (ORFs) in the modified contigs to define candidate genes (**Materials and Methods**). From this composite dataset, we identified 110 pfam domain families that were significantly associated with modified contigs (Bonferroni-corrected *p*-value < 0.01) representing a total of 2365 candidate genes (**Supplementary Table 2**).

The resulting top associations (**Fig. 4a** and **Supplementary Fig. 3a-b**) contained a number of domains families previously found to be involved in DNA synthesis/modification, validating that this approach can uncover relevant functional domains. For example, a subset of enzymes containing the thymidylate synthase domain (PF00303.20) have been shown to produce hydroxymethylpyrimidines (Neuhard et al., 1980). DNA ligase (PF14743.7, PF01068.22), and Cytidine deaminase (PF00383.24) are other domains that have been found in DNA modifying enzymes (Subramanya et al., 1996) (Bhattacharya et al., 1994). In addition, our analysis identified domain families that were not previously known to modify DNA or nucleotides and thus may be novel DNA modifying enzymes or critical regulators.

**Figure 4.**
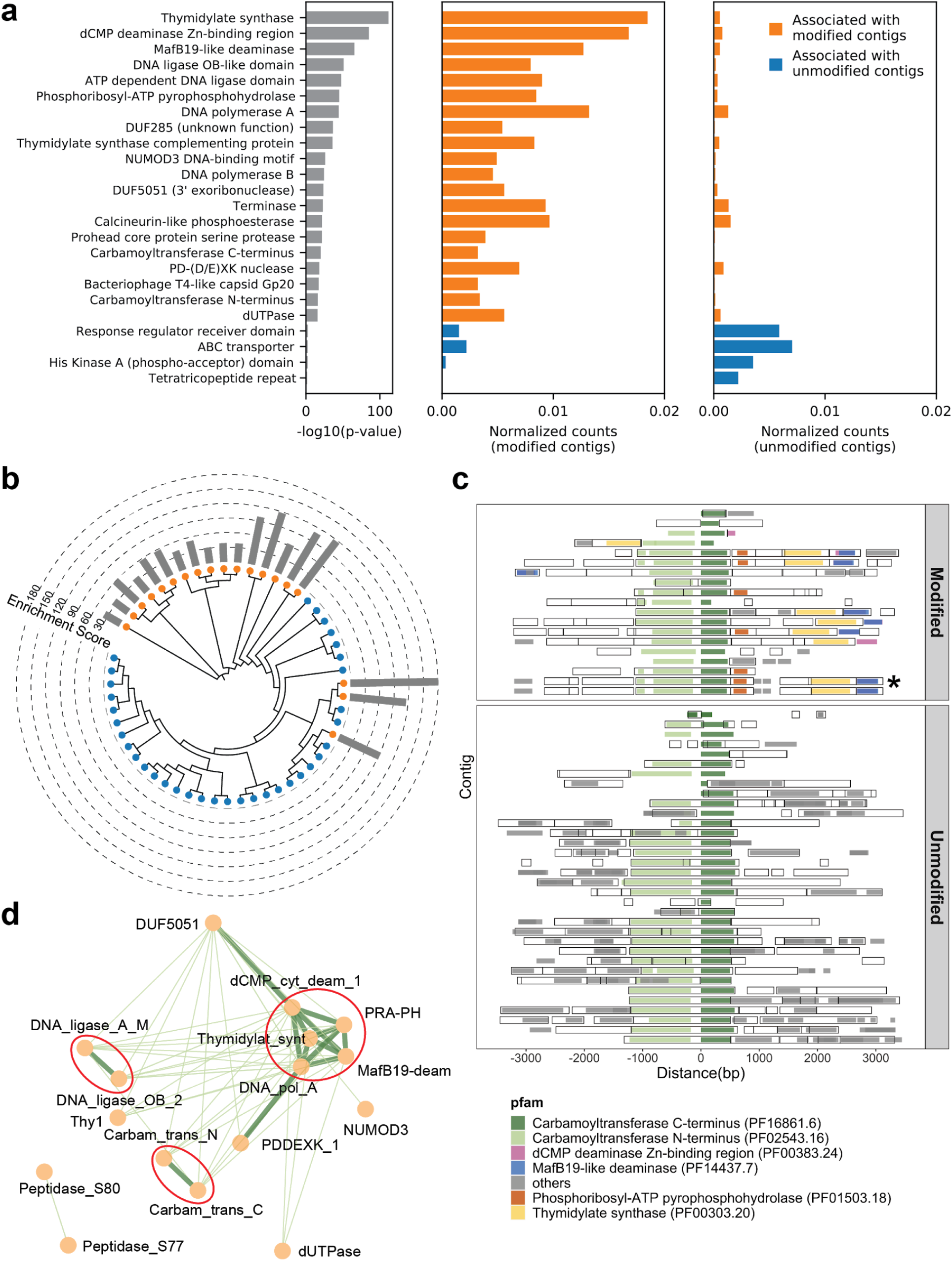
MetaGPA study at domain level. **a**, Identification of associated Pfam protein domains. Left, lists of domains ranked by *p*-values. Middle and right panels represent the normalized counts of each domain for modified or unmodified contigs respectively. The top 20 domains significantly associated with modified contigs were colored in orange and the 4 domains significantly associated with unmodified contigs were colored in blue (Bonferroni-corrected *p*-value <0.01). Data were normalized to total counts in each category (sums of all domain counts in modified or unmodified contigs) respectively. Domains occurring multiple times on the same contigs were counted only once. **b**, Phylogenetic tree of carbamoyltransferase C-terminus domain. Orange and blue dots represent modified and unmodified contigs respectively. Grey bars in the outer ring represent the enrichment scores of each contig with dashed lines showing the scales. Most of the carbamoyltransferase C-terminus domains from modified contigs form a distinctive phylogenetic branch. **c**, Pfam-neighborhood association for carbamoyltransferase C-terminus. Carbamoyltransferase C-terminus is centered in the middle and neighboring Pfams spanning 3 kb upstream and downstream are displayed as solid squares. Colored squares mark the top five Pfams co-occurred with carbamoyltransferase C-terminus in modified contigs. Predicted ORFs on each contig are marked as hollow squares. Asterisk marks the contig containing the enzyme expressed and purified in Fig.6. **d**, Correlation networks between the top 20 associated Pfams with modified contigs. Each node represents a Pfam. The thickness and length of edges were based on the *p*-values between each two nodes with thick and short edges corresponding to more significant relationships (small *p*-values). Only positive correlations with *p*-values < 0.05 are shown. The three interaction cores are circled in red.

To refine the candidate genes, we conducted phylogenetic analysis for each domain family significantly associated with modified contigs. Towards this end, all instances of a particular domain were aligned using a maximum likelihood model and the resulting phylogenetic tree was overlaid with the origin status of the contig (modified/unmodified). Particularly, several domains, including Carbamoyltransferase N-terminus (PF02543.16) and C-terminus (PF16861.6) domains, exhibited a clustered pattern in which most of the sequences from modified contigs formed a distinct phylogenetic clade from the other sequences (**Fig. 4b** and **Supplementary Fig. 3c**). These clades that are almost exclusively derived from modified contigs restating the association of the domain-of-interest with a potential differentiated phenotype of modification. Moreover, this analysis can serve as evidence for refined functional annotation and may suggest a subfamily with specific functions.

Complex DNA modifications in bacteriophages are usually carried out by multiple enzymes whose genes tend to cluster on the genome (Iyer et al., 2013). We therefore extended the analysis to study co-occurrences on the same contigs of domain families associated with modification (**Fig. 4c** and **Supplementary Fig. 3d**). We found for example that the domains most frequently co-occurring with carbamoyltransferase C-terminus (PF16861.6) were carbamoyltransferase N-terminus (PF02543.16), thymidylate synthase (PF00303.20), phosphoribosyl-ATP pyrophosphohydrolase (PF01503.18), dCMP deaminase Zn-binding region (PF00383.24), and MafB19-like deaminase (PF14437.7) (**Fig. 4c**). Congruously, domains belonging to the thymidylate synthase family also co-occurred with carbamoyltransferase N-terminus, phosphoribosyl-ATP pyrophosphohydrolase, dCMP deaminase Zn-binding region, and MafB19-like deaminase domains. Importantly, we found that these co-occurrences are often specific to modified contigs (**Fig. 4c** and **Supplementary Fig. 3d**). For example, the co-occurence of carbamoyltransferase and thymidylate synthase domains is only found in modified contigs (**Fig. 4c**). In unmodified contigs, carbamoyltransferase domains were flanked by genes with unrelated functions such as genes encoding for Glycosyl transferases group 1 or tRNA N6-adenosine threonylcarbamoyltransferase domains.

Together, these results suggested a multi-domain network related to DNA modification. Unbiased co-occurrence analysis was therefore conducted for the top 20 associated domain families. Indeed, significant positive correlations were observed, and the results demonstrated three interaction cores centred on thymidylate synthases, DNA ligases, and carbamoyltransferases respectively (**Fig. 4d**).

#### Differential conservation of amino-acids reveals key residues associated with DNA modification

Having demonstrated that MetaGPA can be successfully deployed to identify protein domains associated with a given phenotype, we next considered whether MetaGPA go as far as identifying associations at residue resolution. To identify such associations, we designed a “differential” conservation score based on existing metrics(Karlin and Brocchieri, 1996; Valdar, 2002) that reflects the degree of association of individual residue with DNA modification (**Materials and Methods**). This score ranks high the residues that show distinct conservation patterns whether they are derived from modified or unmodified DNA. Conversely, weakly conserved residues or residues conserved invariably between modified and unmodified DNA are ranked low.

We used the thymidylate synthase domain alignments to benchmark the differential conservation score. This well characterized family of proteins involved in nucleotide biosynthesis is also ranked by MetaGPA as the most significantly associated with DNA modification (**Fig. 4a**). We aligned the 433 thymidylate synthase protein sequences identified in our composite dataset with the canonical *E. coli* thymidylate synthase which has been structurally and biochemically characterized (Stout et al., 1998a). Our differential conservation score identifies N177 and H147 of *E. coli* thymidylate synthase as the top 2 positions (**Fig. 5a**). Both residues are within the enzyme’s active site (**Fig. 5c**) (Stout et al., 1998b) and have been substituted for the most part with D and T in the thymidylate synthases found in modified contigs (**Fig. 5b**). N/D substitution at N177, has previously been identified as the key substitution responsible for changing the enzyme substrate specificity from uracil (N residue) to cytosine (D residue) (Hardy and Nalivaika, 1992) (Graves et al., 1992) (Liu and Santi, 1992). The H147 forms a hydrogen bonding network through a water molecule to the keto oxygen of position 4 in dUMP (Matthews et al., 1990), suggesting a role in substrate specificity. Alternatively, H147 is part of a H-bond network with Y94, a key residue important for proton abstraction at C5. Therefore, residue occupancy at position 147 may accommodate differences between the pKa of C5 between cytosine and uracil substrates (Hong et al., 2007). Taken together, our differential conservation score identified functionally critical residues for thymidylate synthase activity and substrate specificity.

**Figure 5:**
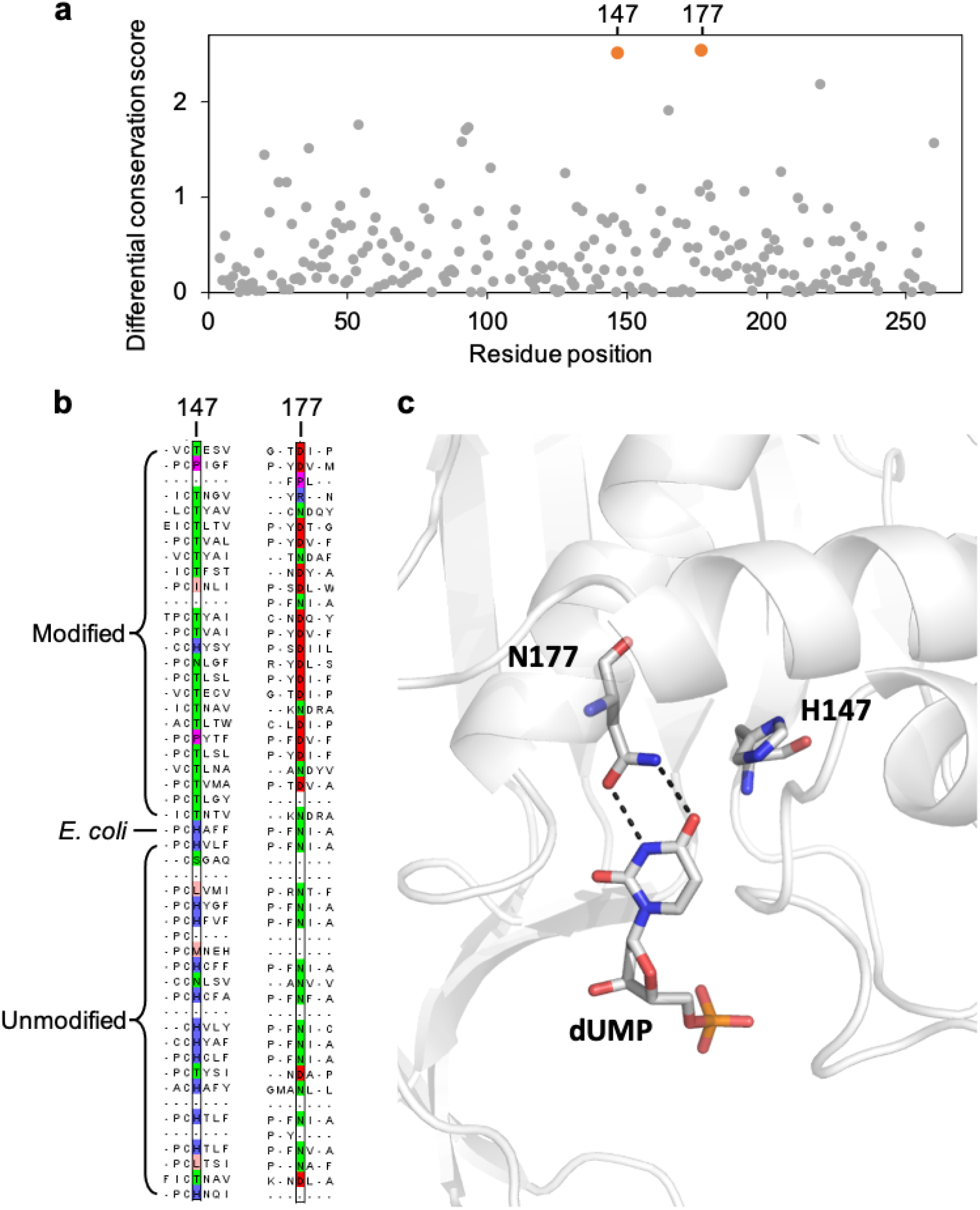
Key residues in the thymidylate synthase associated with DNA modification. **a**, Differential conservation score for each position in the thymidylate synthase alignment. Positions are relative to the *E. coli* thymidylate synthase. **b**, Multiple sequence alignment of 25 and 24 randomly selected thymidylate synthase sequences from the modified and unmodified contigs respectively together with the *E. coli* thymidylate synthase. Aligned residues at positions 147 and 177 (relative to *E. coli* thymidylate synthase) are colored according to the physicochemical properties of the amino acids. **c**, Structure of the active site of *E. coli* thymidylate synthase (PDB 1BID) highlighting the H147 and N177 residues and the dUMP substrate.

We then performed the same analysis with the top 20 associated Pfam domains from our list (**Fig. 4a**). Among them, carbamoyltransferase sequences were aligned with the O-carbamoyltransferase family member TobZ that has been structurally and functionally characterized (Parthier et al., 2012). Our analysis identified three residues (W408, G421 and R423) with the highest differential conservation scores (**Supplementary Fig. 4a-b**). All three residues are located within 14 Å to the carbamoyl phosphate and ADP binding pocket and may be important in defining the enzyme’s specificity towards cytosine derivatives (**Supplementary Fig. 4c**).

Altogether, these results indicate that MetaGPA is able to associate single residue within a protein to the studied phenotype.

### 3. Characterization of a novel modifying enzyme – 5-hydroxymethylcytosine carbamoyltransferase

Among the 110 domains that are significantly associated with DNA modifications, carbamoyltransferases are ranking high in our MetaGPA study and are exhibiting all three of our refinement metrics. Carbamoyltransferases are part of a large protein family catalyzing the addition of a carbamoyl group to various substrates but so far none of them has been shown to act on any form of cytosine, potentially revealing a new function and a new cytosine modification. We therefore sought to further explore these enzymes for their ability to modify cytosine.

The co-occurrence of carbamoyltransferase and thymidylate synthase homologs specifically in modified contigs (**Fig. 4c**) resembles the arrangement of the T4 phages for which genes coding for the dCMP hydroxymethylase (a member of the thymidylate synthase family) and β-glucosyltransferase co-occur on the genome (Miller et al., 2003). As for the products of those genes, the T4 dCMP hydroxymethylase catalyses the formation of 5hmdC, while β-glucosyltransferase transfers a glucose moiety to the 5hmdC residues. This analogy led us to hypothesize a novel form of DNA modification, in which the carbamoyltransferase catalyzes the reaction from 5hmdC to 5-carbamoyloxymethylcytosine (5cmdC) (**Fig. 6a**).

**Figure 6.**
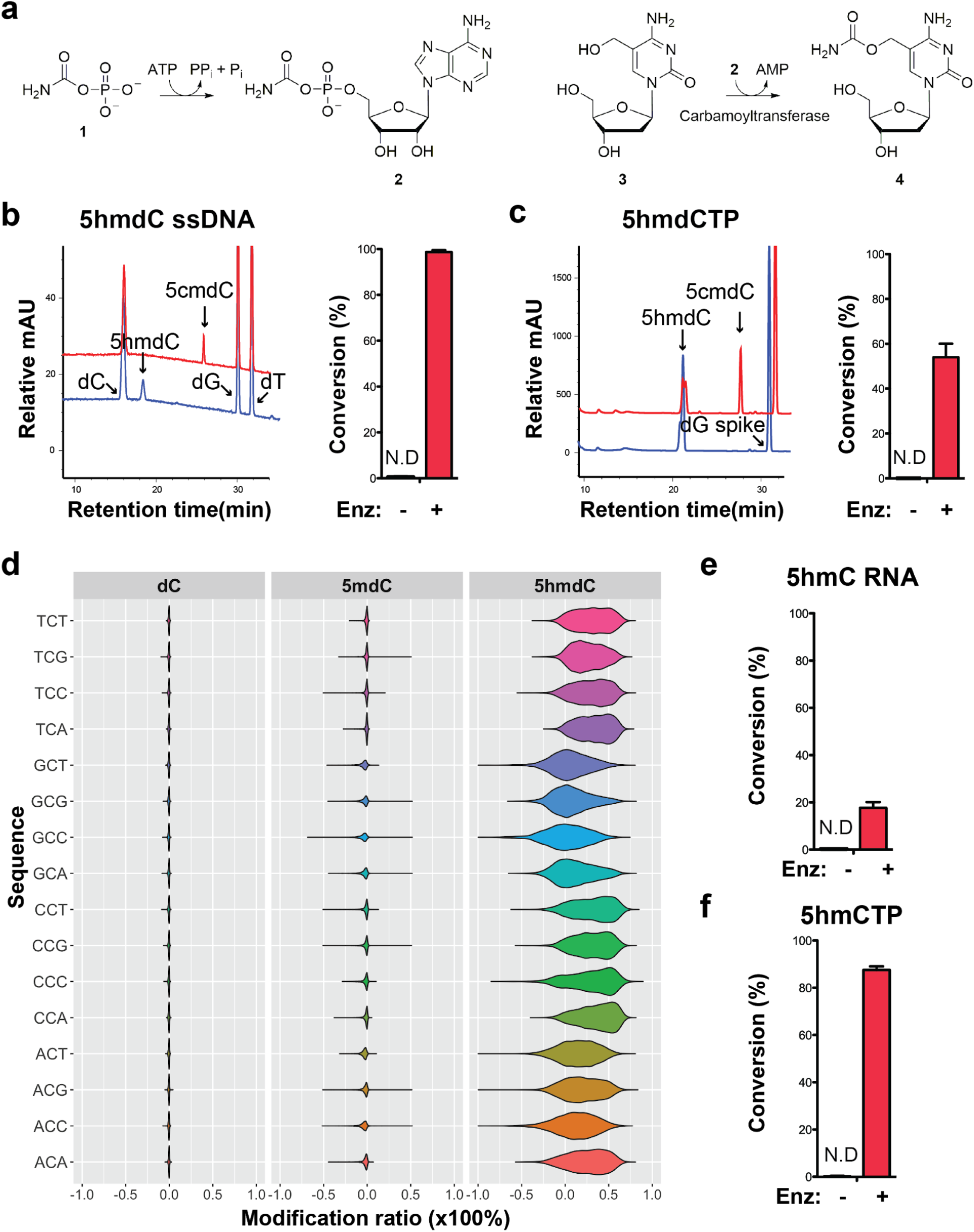
Validation of the novel 5-hydroxymethylcytosine carbamoyltransferase. **a**, Predicted reactions. **1**, carbamoylphosphate; **2**, carbamoyladenylate intermediate; **3**, 5-hydroxymethylcytidine; **4**, 5-carbamoyloxymethylcytidine. **b**, LC-MS trace and quantification for enzymatic reaction with a single-stranded DNA oligo containing an internal 5hmdC. Bar graph represents average conversion rates ± SEM from three independent experiments using three different DNA oligos. N.D., not detected. **c**, LC-MS trace and quantification for enzymatic reaction with 5hmdCTP nucleoside triphosphate. Bar graph shows average conversion rates ± SEM from three independent experiments. **d**, Enzyme preference on NCN sequences. Genomic DNA from lambda (dC), XP12 (5mdC) and T4gt (5hmdC) were mixed at 1:1:1 ratio by molarity before being subjected to enzymatic selection. Modified ratio of each C in NCN motifs (with N being A, T, C, or G) was normalized to the no-enzyme control. **e**, LC-MS quantification for enzymatic reaction with a single-stranded RNA oligo containing an internal 5hmC. Bar graph shows average conversion rates ± SEM from three independent experiments. **f**, Quantification for enzymatic reaction with 5hmCTP. Bar graph shows average conversion rates ± SEM from three independent experiments. Oligonucleotides and nucleoside triphosphate were converted to nucleosides before LC-MS analysis. A dG spike was added as internal reference for quantification of nucleoside triphosphates.

Our composite dataset contains 62 genes annotated as carbamoyltransferase for which 17 are found in the modified contigs. We selected a carbamoyltransferase candidate gene from a modified contig originally sequenced in sewage #2 sample containing both the thymidylate synthase and the carbamoyltransferase genes (**Supplementary Fig. 5a**). The ORF was cloned into pET28b vector, and the 63 kDa protein product was expressed in *E.coli* and purified (**Supplementary Fig. 5b**). The predicted reaction was tested by enzymatic assays and results showed that every substrate, namely carbamoyl phosphate, ATP, 5hmdC (genomic T4gt DNA was used as substrate in these experiments) and the enzyme were indispensable for the reaction (**Supplementary Fig. 5c-d**). The expected product was detected by liquid chromatography-mass spectrometry (LC-MS) and confirmed with corresponding fragmentation patterns (**Supplementary Fig. 5e**). Nearly 70% of 5hmdC were converted into 5cmdC in the denatured single-stranded T4gt genomic DNA. Interestingly, the activity of our carbamoyltransferase was several fold lower on double-stranded DNA, suggesting the preference of this enzyme for single-stranded DNA (**Supplementary Fig. 5c**). When using synthesized single-stranded DNA oligo containing a single internal 5hmdC site as substrate, the conversion rate was nearly 100% (**Fig. 6b**). We also tested if the carbamoyltransferase could react with free deoxynucleoside triphosphate. LC-MS results demonstrated about 60% conversion of 5-hydroxymethyl-2’-deoxycytidine triphosphate (5hmdCTP) (**Fig. 6c**). As expected, no activity was seen for 5-methyl-2’-deoxycytidine triphosphate (5mdCTP) or 5-hydroxymethyl-2’-deoxyuridine triphosphate (5hmdUTP), indicating the carbamoyltransferase is specific to 5hmdCTP. The fact that the carbamoyltransferase is active on 5hmdCTP (**Fig. 6c** and **Supplementary Fig. 5f**) opens up the possibility that the reaction could take place before the nucleotide is incorporated into the phage DNA.

To examine if the carbamoyltransferase favors certain DNA sequences, we performed the enzymatic assay on a mixed genomic DNA library containing Lambda (dC), XP12 (5mdC) and T4gt (5hmdC). Both untreated (control) and treated libraries were subjected to APOBEC3A deamination (see **Materials and Methods**). Carbamoylation protects cytosine derivatives from deamination by APOBEC and thus, the difference in deamination rate between control and treated libraries is indicative of carbamoylation. We saw a decrease in deamination rate only in the T4gt genome indicating specific carbamoylation on 5hmdC. This result further validates the LC-MS results regarding the specificity of the enzyme (**Fig. 6d**). Furthermore, all combinations of NCN motifs containing 5hmdC displayed comparable deamination levels compared to the control library, suggesting a general binding mechanism with no preferred context. This result was also consistent with enzymatic assays performed on free nucleotides, which further suggests that the *in-vivo* carbamoylation reaction may take place prior to DNA replication.

Based on our finding that the carbamoyltransferase prefers single-stranded DNA, we further investigated whether RNA containing 5-hydroxymethylcytosine can be modified by the enzyme. LC-MS and fragmentation pattern confirmed that 5cmC is formed during the reaction, albeit at much lower yields (**Fig. 6e** and **Supplementary Fig. 5g-h**).

Carbamoylation of free nucleotide 5-hydroxymethylcytidine triphosphate (5hmCTP) was also detected (**Fig. 6f** and **Supplementary Fig. 5f**). We thus concluded that this novel nucleic acid modifying enzyme identified from MetaGPA studies acts on DNA, RNA and nucleotide triphosphates.

## Discussion

In this work, we report a novel analytic framework called MetaGPA that links functional phenotype with genetic information in metagenomes. Being cost effective and requiring reduced efforts, MetaGPA essentially rests on two sequencing libraries derived from the same starting material: one library reflects all the organisms in the community while the other library results from a selection to only contain organisms with a phenotype of interest. This experimental setup allows for candidate genes identification in less than a week.

Our MetaGPA study framework is phenotype-driven and hypothesis-free. Theoretically, it can be performed with any case/control cohort pair as long as distinct phenotypes can be partitioned through selection of the case cohort. For example, these phenotypes may include but are not limited to DNA modifications, phage sensitivity to chemicals and cell surface determinants such as O-antigen. Nonetheless, this partitioning is crucial for MetaGPA to succeed and therefore requires the development and optimization of the selection process for every new MetGPA application.

We used this approach to screen environmental phage metagenomes and successfully identified 110 domains representing 2365 candidate proteins, mostly enzymes that are significantly associated with DNA modifications. From these candidates, we validated a novel DNA/RNA modifying enzyme responsible for the previously unknown 5-carbamoyloxymethylcytosine modification. To our knowledge, the only reported carbamoylated nucleotide is the 5-carbamoylmethyluridine, which is located in the first position of the anticodon of yeast valine tRNA34 (Yamamoto et al., 1985). Importantly, we predicted, at single residue resolution, the key positions in the enzyme associated with this DNA modification, enabling candidate prioritisation and protein engineering.

In these MetaGPA experiments, the sequencing libraries can be directly used as material for selection because the phenotype of interest (cytosine modification) is covalently attached to the genetic material. However, for other scenarios for which phenotypes are not physically coupled with DNA or RNA, selection will have to be done while preserving the integrity of cells and viral particles to retain the link between phenotype and genotype.

MetaGPA provides notable benefit for *de novo* exploratory discoveries especially with largely unknown microbial communities such as the virulome where existing knowledge is limited. Besides the example provided in this study, we expect applications of our framework to be expanded to a wide range of topics, e.g. antibiotic resistance, microbial responses to atypical environments, and host-microbe interaction.

## Acknowledgement

The authors would like to thank Barry Cohen and the junior team (Grace, Alina, Camille and Jeanne) for environment sample collections, Lana Saleh for useful discussions and providing the RNA oligos, Vladimir Potapov, Zhiyi Sun and Tamas Vincze for analysis and IT support and the sequencing core. John Buswell for providing the Pyrrolo dC adaptor. This work was supported by New England Biolabs.

## Data access

All raw and processed sequencing data generated in this study have been submitted to the NCBI Sequence Read Archive (SRA; https://www.ncbi.nlm.nih.gov/sra) under accession number PRJNA714147.

## Materials and Methods

### Genomic DNA

The E. coli K-12 MG1655, XP12 and T4gt genomic DNA used in this study were obtained from NEB.

### Environmental phage collection

For each batch, 2 ~ 4 liters of sewage or coastal seawater were used for phage collection. Large debris and bacterial cells were pelleted and removed by centrifuging at 5,000 x*g* for 30 min at 4 °C. Phage particles in the supernatant were precipitated by adding PEG8000 to 10% (w/v) and NaCl to 1 M and let stand at 4 °C overnight. Aggregates of phage particles were pelleted at 10,000 x*g* for 30 min at 4 °C, washed with 1 mL solution containing 10% PEG8000 and 1 M NaCl, and resuspended in 2 ~ 4 mL of phage dilution buffer (10 mM Tris-HCl at pH 8.0, 10 mM MgCl_2_, 75 mM NaCl). The crude phage particle suspension was stored at 4 °C for subsequent phenol-chloroform DNA extraction.

### Phenol-chloroform DNA extraction

2 ~ 4 mL of crude phage suspension was divided in 400 μL aliquots. For each aliquot, phage particles were lysed at 56 °C for 2 h in 550 μL of lysis buffer (100 mM Tris-HCl at pH 8.0, 27.3 mM EDTA, 2% SDS, ~1.6 U Proteinase K [NEB #P8107]). After lysis, RNase A was added to 10 μg/mL and the reaction was incubated at 37 °C for 30 min. 1X volume (~550 μL) of phenol-chloroform (Tris-HCl buffered at pH 8.0) was mixed with the lysis solution and vortexed vigorously for ~1 min, and centrifuged at 10,000 *xg* for 5 min for phase separation. The top aqueous layer (~500 μL) was collected and mixed with 1X volume (~500 μL) of chloroform, vortex vigorously, and centrifuged for phase separation. The top aqueous layer (~450 μL) was collected. 1X volume of isopropanol was slowly added on top of the aqueous solution. Phage DNA was “spooled” with a glass capillary by swirling and mixing isopropanol with the aqueous solution. The spooled DNA was washed in 70% ethanol, dried at room temperature for ~30 min, and dissolved in ~600-800 μL of TE buffer (10 mM Tris pH 7.5, 1 mM EDTA).

The phage DNA solution was further purified by ethanol precipitation. Briefly, DNA was precipitated by adding 0.1X volume of 3 M sodium acetate and 2.5X volume of 100% ethanol and incubated at −20 °C overnight. Precipitated DNA was pelleted at 16,000 x*g* for 20 min, washed twice with 1 mL of 70% ethanol, dried at room temperature, and finally dissolved in 200 μL of TE buffer for storage at −20 °C. On average more than 20 μg of DNA was extracted in each batch.

### Illumina library preparation

For each environmental sample, 1 μg of metagenomic DNA was sheared to 300 bp in 130 μL of TE buffer (10 mM Tris pH 7.5, 1 mM EDTA) using Covaris S2 Focused Ultrasonicator. 1.3 μL of 10 mg/mL RNase A (Qiagen #1007885) was added and incubated at 37 °C for 30 min to remove RNA. To remove EDTA, the sheared DNA was purified with Zymo Oligo Clean & Concentrator™ Kit (#D4061) and eluted in 50 μL of 1 mM Tris buffer (pH 7.5).

One reaction of NEBNext^®^ Ultra™ II DNA Library Prep Kit for Illumina^®^ (NEB #E7645) was used for 1 μg of input DNA, with the following modifications to the standard protocol: 5mdC Y-shaped Illumina adaptors were used to protect the adaptor from subsequent enzymatic treatment; the dCTP was replaced with 5mdCTP in the end repair reaction. The DNA library was purified with 1X volume of NEBNext^®^ Sample Purification Beads (NEB #E7103) and eluted with 40 μL of 1 mM Tris buffer (pH 7.5).

For each of the two sewage DNA samples, experiments were performed in duplicate. each one contained two pairs of replicate libraries subjected to enzymatic selection or control respectively, The coastal sample generated only one pair: one library for enzymatic selection and one for control.

### Enzymatic selection protocol

First to test the recovery of modified DNA with enzymatic selection, mixed genomic DNA (E. coli and T4gt) were prepared at various dilutions. A total of 250 ng mixed DNA was used per reaction. 1 μL TET2 (NEB #7120S) and 1 μL T4-BGT (NEB #M0357S) were added to the 50 μL reaction mixture containing 1x TET2 reaction buffer, 40 μM UDP-Glucose and 40 μM iron(ii) sulfate hexahydrate. After 60 min incubation at 37 °C, Proteinase K was added at 0.4 mg/mL to inactivate the enzymes. Products were purified with Zymo Oligo Clean & Concentrator Kit (#D4061) and eluted in 16 μL water. To denature double-stranded DNA, 4 μL formamide (Sigma #11814320001) was added. The 20 μL mixture was then incubated at 95 °C for 10 min and immediately transferred to an ice bath. One μL APOBEC3A (NEB #E7120S) was added directly to the reaction with 10 μL of 10x APOBEC3A reaction buffer and 1 μL BSA (10 mg/mL). The reaction volume was brought up to 100 μL with water. APOBEC3A-mediated deamination was conducted at 37 °C for 3 h. Purification was performed using Zymo Oligo Clean & Concentrator kit and elution with 43 μL of water. In the final step, the reaction was incubated with 2 μL of USER (NEB #5508) in 1X CutSmart Buffer at 37 °C for 15 min before final purification with Zymo Oligo Clean & Concentrator kit. Purified samples were then used for quantitative PCR in the next step.

For each prepared phage library sample, 100 ng spiked-in genomic DNA mixture (*E. coli*:XP12:T4gt = 1:1:1 by molarity) were added to the library before being subjected to enzymatic selection protocol described above. The final library was eluted in 50 μL of 1 mM Tris buffer (pH 7.5).

### Quantitative PCR

The qPCR reactions were performed with enzymatic selection or control samples using Luna Universal qPCR Master Mix (NEB #M3003S) on a Bio-Rad CFX96 real-time PCR detection system. Two μL of purified DNA (diluted 100 folds) were added per reaction. Primers used in the experiments were the following: *E. coli* F: 5’-TTGCTGAGTTTCACGCTTGC, *E. coli* R: 5’-AAAACCGCTTGTGGATTGCC, T4gt F: 5’-TCGCGAAACGGTTTTCCAAG, T4gt R: 5’-AAAGCGCTTGACCCAACAAC, XP12 F: 5’-TGCGATGTTGGATTCGTTGG, and XP12 R: 5’-ACAACCCGCCATAATGGAAC. Recovery was normalized to control using the delta-delta Ct method.

### Illumina sequencing

Libraries were indexed (with NEBNext^®^ Multiplex Oligos for Illumina E7335), amplified using NEBNext^®^ Ultra™ II Q5^®^ Master Mix (6 cycles for control library and 12 cycles for selection library) and pooled for sequencing on an Illumina Nextseq instrument with paired end reads of 75bp.

### Sequencing data processing

Paired-end reads were downloaded as FASTQ files and trimmed with Trim Galore v0.6.4 (https://www.bioinformatics.babraham.ac.uk/projects/trim_galore/) using the --paired option. K-mer counting from reads was done with JELLYFISH v2.2.10 and 16-mer was chosen based on best resolution. *De novo* assembly of contigs for each sample was performed with SPAdes v3.13.0 with the --meta option. We selectively reported contigs longer or equal to 1000 bp. To remove redundant contigs between selection and control pairs from each experiment, we used CD-HIT v4.8.1 nucleotide mode cd-hit-est with sequence identity threshold set to 0.95. Other options used were -n 10 -d 0 -M 0 -T 4. The remaining non-redundant contigs were annotated with HMM-based Pfam entries (Pfam-A) using HMMER v3.3. Mapping of reads onto contigs was done with BOWTIE2 v2.3.5.1 together with SAMTOOLS v1.9 to generate, sort and index bam files for later analysis. ORFs on each contig were predicted with GLIMMER 3.02. We used the long-orfs program with options --cutoff 1.15 and --linear to identify long orfs that are very likely to contain genes. Other options used in glimmer3 program were --max_olap 100, --gene_len 110 and --threshold 30.

### Contig enrichment score calculation

The enrichment score for each contig was calculated using the normalized mapped reads (reads per kb per million, RPKM) from selection and control as follows: enrichment score = RPKM_(selection)_ / RPKM_(control)_. The mapped reads counts were generated with Multicov using BEDTOOLS v2.29.2. Contigs with higher enrichment score represent more mapped reads in selection library relative to control library, therefore, are more likely to be associated with modification. We considered contigs with an enrichment score greater or equal to 3 to be modified and the rest unmodified. The calculation was done individually for three independent experiments.

### Fisher’s exact test and correction

The information including the number and type of Pfams on each contig was obtained with hmmsearch in the annotation step. We then re-organized the data and counted the number of contigs containing each type of Pfam in the control or the selection group. To avoid redundant counting, Pfams occurred multiple times on the same contig was counted only once. Fisher’s exact test was performed for each Pfam to identify if the count difference between the selection and control group is significant. Because large-scale multiple testing was conducted for each Pfam, we did the Bonferroni correction to adjust the *p*-value. Both tests were performed in python with SciPy or Statsmodels modules.

### Phylogenetic analysis

For each Pfam of interest, the protein sequences from contigs containing the Pfam were aligned with MUSCLE v3.8.1551. The resulting aligned fasta files were subjected to construct phylogenetic trees using the maximum likelihood method in the phylogenetic analysis program RAxML v8.2.12. We chose the -f a option to do rapid bootstrap analysis and the -m PROTGAMMAAUTO model to automatically determine the best protein substitution model to be used for the dataset. The parsimony trees were built with random seeds 1237. The online tool iTOL (https://itol.embl.de/) was used to visualize trees.

### Co-occurrence network analysis

The presence-absence matrix with rows being the Pfams and columns being the contigs was generated with annotation output file from the previous step. We specifically performed co-occurrence analysis in the R package coocur v1.3 for the top 20 Pfams associated with modified contigs. Significant positive correlations (*p*-value < 0.05) were exported and the network was visualized in Cytoscape v3.8.0 with prefuse force directed layout.

### Differential conservation score

Protein sequences were assigned to two groups according to whether they were encoded on modified or unmodified DNA. After multiple sequence alignment, positions that have less than 50% residues present were ignored. Differential conservation score was calculated at each aligned position. For each position in the alignment, intra-group similarity scores were calculated by the average of all possible “within-group” pairwise similarities, while the inter-group similarity score was calculated from all possible “across-group” pairwise similarities using the BLOSUM80 matrix. For a given multiple sequence alignment column, let *N*_1_ and *N*_2_ be the number of residues for the modified and unmodified groups, respectively, the two intra-group similarity scores (I_modified_ and I_unmodified_) were defined as

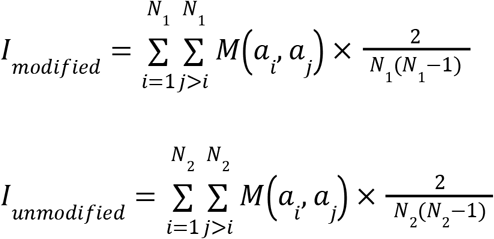

where *M*(*a_i_, a_j_*) is the value of amino acid pair *a_i_* and *a_j_* in the BLOSUM80 matrix. The inter-group similarity score (J) was defined as

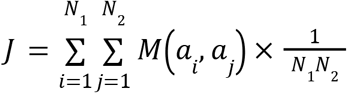

The differential conservation score (S) was defined as the average of two intra-group similarity scores subtracted by the inter-group similarity score.

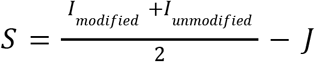

### Expression and purification of carbamoyltransferase

The carbamoyltransferase sequence was extracted from *de novo* assembled contigs. The expression plasmid was synthesized from GenScript. Two 6x His-tags were co-expressed at both the N-terminus and the C-terminus of the recombinant protein using T7 Express Competent *E. coli* (NEB #C2566). Cells were cultured in LB media until an OD600 of 0.6 and induced with 0.4 mM IPTG (Growcells #MESP-2002) for protein expression. One μM Iron (II) was also added to facilitate protein folding. The induced cultures were maintained at 16 °C in a shaker at 200 rpm for 23 h. Cells were harvested by spinning down cell pellets at 3500 rpm at 4 °C for 30 min. Cell pellets from 4 L culture were resuspended in 160 mL buffer A containing 20 mM Tris pH 7.5, 500 mM NaCl, 0.05% Tween-20, 20 mM imidazole and sonicated using a Misonix S-4000 sonicator with 20 s on and 20 s off cycles until an OD260 plateau was reached. Cell lysates were spinned down at 13,000 rpm for 30 min in a pre-chilled centrifuge at 4 °C. The supernatant was separated and combined with 0.2 mM PMSF (Sigma #78830). 50 mL of supernatant was loaded on AKTA (GE Healthcare Life Sciences) with 1 mL Histrap column (GE Healthcare Life Sciences) pre-equilibrated with buffer A. The column was washed with 50 mL buffer A and eluted with a gradient of buffer B containing 20 mM Tris pH 7.5, 500 mM NaCl, 0.05% Tween-20, and 500 mM imidazole. Aliquots containing concentrated proteins were pooled and diluted 1:1 with 20 mM Tris pH 7.5, 5% glycerol and 0.05% Tween-20. The diluent was reloaded on AKTA with 5 mL Hitrap Q HP column (GE Healthcare Life Sciences), followed by a wash with 35 mL buffer containing 20 mM Tris pH 7.5, 100 mM NaCl, 5% glycerol, and 0.05% Tween-20 and eluted with a gradient of buffer containing 20 mM Tris pH 7.5, 1 M NaCl, 5% glycerol, and 0.05% Tween-20. Finally, collected fractions with concentrated proteins were pooled and mixed with equal volume glycerol for storage at −20 °C.

### Carbamoyltransferase enzyme assay

For enzyme assay using T4gt genomic DNA as substrate, 10 min incubation at 95 °C was performed to denature double-stranded DNA and the sample was immediately transferred to ice bath to prevent re-annealing. Then 0.38 nM denatured DNA was used for each 50 μL reaction with 1x NEBBuffer2.1 (NEB #B7202S), freshly prepared 10 μM Iron(II) sulfate hexahydrate (Sigma #203505), freshly prepared 10 mM carbamoylphosphate and 5 mM ATP. Carbamoyltranferase was added to the reaction at 7.2 μM. The reaction mixture was incubated at 30 °C for 3 h before adding 2 μL Proteinase K to inactivate the enzyme. After 30 min incubation at 37 °C with Proteinase K, DNA was purified with Zymo Oligo Clean & Concentrator kit. For assays with synthesized single-stranded DNA oligos containing 5hmdC, the heat-denaturing step was omitted. Oligos were added at 1.6 μM per 50 μl reaction with the same concentration of carbamoyltransferase and other components added as listed before. Purification was performed using Norgenbiotek Oligo Clean-up and Concentrator kit (#34100). For assays with free nucleotides, 0.5 mM of the corresponding nucleotide was used per reaction. For assays with synthesized RNA oligos containing 5hmC, 1.57 μM RNA was added per reaction.

### LC-MS and fragmentation analysis

Genomic DNA and synthetic oligonucleotides were digested to nucleosides by treatment with the Nucleoside Digestion Mix (NEB #M0649S) at 37 °C for 3 h. The resulting nucleoside mixtures were directly analyzed by reversed-phase LC/MS or LC-MS/MS without further purification. Nucleoside and Nucleotide analyses were performed on an Agilent LC/MS System 1200 Series instrument equipped with a G1315D diode array detector and a 6120 Single Quadrupole Mass Detector operating in positive (+ESI) and negative (-ESI) electrospray ionization modes. LC was carried out on a Waters Atlantis T3 column (4.6 mm × 150 mm, 3 μm) at a flow rate of 0.5 mL/min with a gradient mobile phase consisting of 10 mM aqueous ammonium acetate (pH 4.5) and methanol. MS data acquisition was recorded in total ion chromatogram (TIC) mode. LC-MS/MS was performed on an Agilent 1290 UHPLC equipped with a G4212A diode array detector and a 6490A triple quadrupole mass detector operating in the positive electrospray ionization mode (+ESI). UHPLC was performed on a Waters XSelect HSS T3 XP column (2.1 × 100 mm, 2.5 μm particle size) at a flow rate of 0.6 mL/min with a binary with a gradient mobile phase consisting of 10 mM aqueous ammonium formate (pH 4.4) and methanol. MS/MS fragmentation spectra were obtained by collision-induced dissociation (CID) in the positive product ion mode with the following parameters: gas temperature 230 °C, gas flow 13 L/min, nebulizer 40 psi, sheath gas temperature 400 °C, sheath gas flow 12 L/min, capillary voltage 3 kV, nozzle voltage 0 kV, and collision energy 5-65 V.

### Sequence preference of carbamoyltransferase

Library preparation was performed except the following modifications to the standard protocol: 1) we did not perform RNase A treatment for this experiment; 2) we used pyrrolo-dC Y-shaped adaptor instead of regular adaptor so that they are protected from subsequent enzymatic treatment. For each library, 1 μg genomic DNA mixture (Lambda:XP12:T4gt = 1:1:1 by molarity) was used. After adapter ligation, DNA libraries were purified with 1X volume of NEBNext^®^ Sample Purification Beads (NEB #E7103) and eluted with 20 μL nuclease free water. The sample was then denatured by heating to 95 °C for 10min and subjected to carbamoyltransferase reaction as described above. Carbamoyltranferase was added to the reaction at 7.2 μM for every 1ug DNA library. Purification was performed using Zymo Oligo Clean & Concentrator kit and eluted with 16 μL water. Purified DNA samples were heated at 90 °C with 4 μL formamide to generate single-stranded fragments for the deamination reaction. One μL APOBEC3A was added per reaction to both carbamoyltransferase-treated or control (untreated) samples with 10 μL of 10x APOBEC3A reaction buffer and 1 μL BSA (10 mg/mL). The reaction mixture was incubated at 37 °C overnight. Final libraries were purified using Zymo Clean & Concentrator kit, indexed (with NEBNext^®^ Multiplex Oligos for Illumina E7335) and amplified with NEBNext Q5U Mater mix(NEB #M0597). Sequencing was performed on an Illumina Mi-seq instrument with pair-end reads (2×75bp). Raw reads were trimmed with TrimGalore. Methylation was analyzed with Bismark v0.22.3 and plotted with RStudio v3.6.3.

### Synthesis of 5hmC RNA oligonucleotide

Forward and reverse DNA templates were annealed at 95 °C for 4 min and slowly cooled for 20 min. RNA synthesis was performed with HiScribe T7 High Yield RNA Synthesis Kit (NEB #E2040). One μg of annealed DNA template was used per reaction with 1.5 μL T7 RNA Polymerase Mix. 5hmCTP was used with the other three nucleotides ATP, UTP and GTP at 7.5 mM each. The reaction was incubated at 37 °C for 4 h. Two μL Nulease-free DNase I were added to each reaction to digest DNA templates, followed by incubation at 37 °C for 15 min. Synthesized RNA was purified with Norgenbiotek Oligo and Concentrator kit and stored at −80°C.

### Nucleotides and synthesized oligos

Single-stranded DNA oligos used in enzymatic assays were purchased from IDT. The sequences are as follows:

5hmdC-1: 5’-TGTCCGATAGACT{5hmdC}TACGCA;

5hmdC-2: 5’-AACTCGCCGAGGATTT{5hmdC}TAC;

5hmdC-3: 5’-{Fam-AmC6}ACACCCATCACATTTACAC{5hmdC}GGGAAAGAGTTGAATGTAGAGTTGG.

The DNA templates for synthesizing RNA were purchased from IDT as follows (T7 promoter sequence was underlined):

Forward: 5’-<u>GACCTAATACGACTCACTATAG</u>GGAGTGAGAAGATGGTCTAGGTGTTTATTGGTGATGAA ComRev: 5’-TTCATCACCAATAAACACCTAGACCATCTTCTCACTCC<u>CTATAGTGAGTCGTATTAGGTC</u>. 5hmdCTP (D1045) and 5mdCTP (D1035) were purchased from Zymo Research. 5hmdUTP (N-2059) and 5hmCTP (N-1087) were purchased from Trilink Biotechnologies.

## Code availability

Custom-built bioinformatics pipelines are available at https://github.com/linyc74/MetaGPA

