## Supplementary Figures and Tables for "A Genome-Phenome Association study in native microbiomes identifies a mechanism for cytosine modification in DNA and RNA"

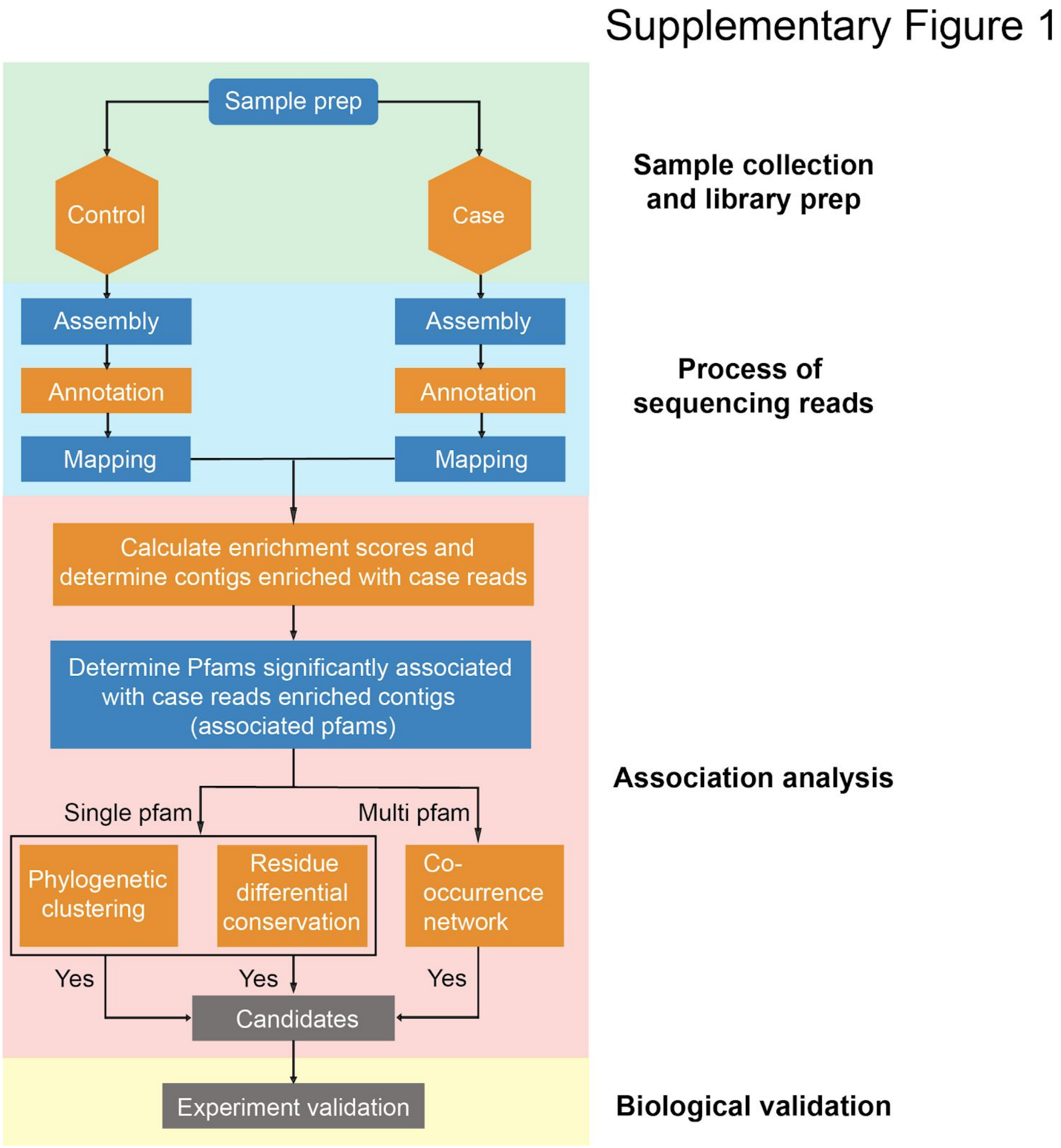


**Supplementary Figure 1** Detailed workflow of MetaGPA studies. Environmental samples are subjected (case) or not (control) to a defined selection to retain only the genetic material from microorganisms that harbor the desired phenotype. Control and case cohorts are sequenced using Illumina and the resulting reads are assembled and mapped. Annotation is performed on predicted genes using HMMER and the Pfam protein domain database. Next, protein domain families that are significantly associated with the studied phenotype are identified. For each significant domain family, the respective genes found in the case (candidate genes) and control cohorts are extracted and subjected to phylogenetic and residue differential conservation analysis. Finally the co-occurence of multiple candidate genes within the same contig reveals biochemical pathways. Together, these metrics prioritized candidate genes for further experimental validation.

### Supplementary Figure 2

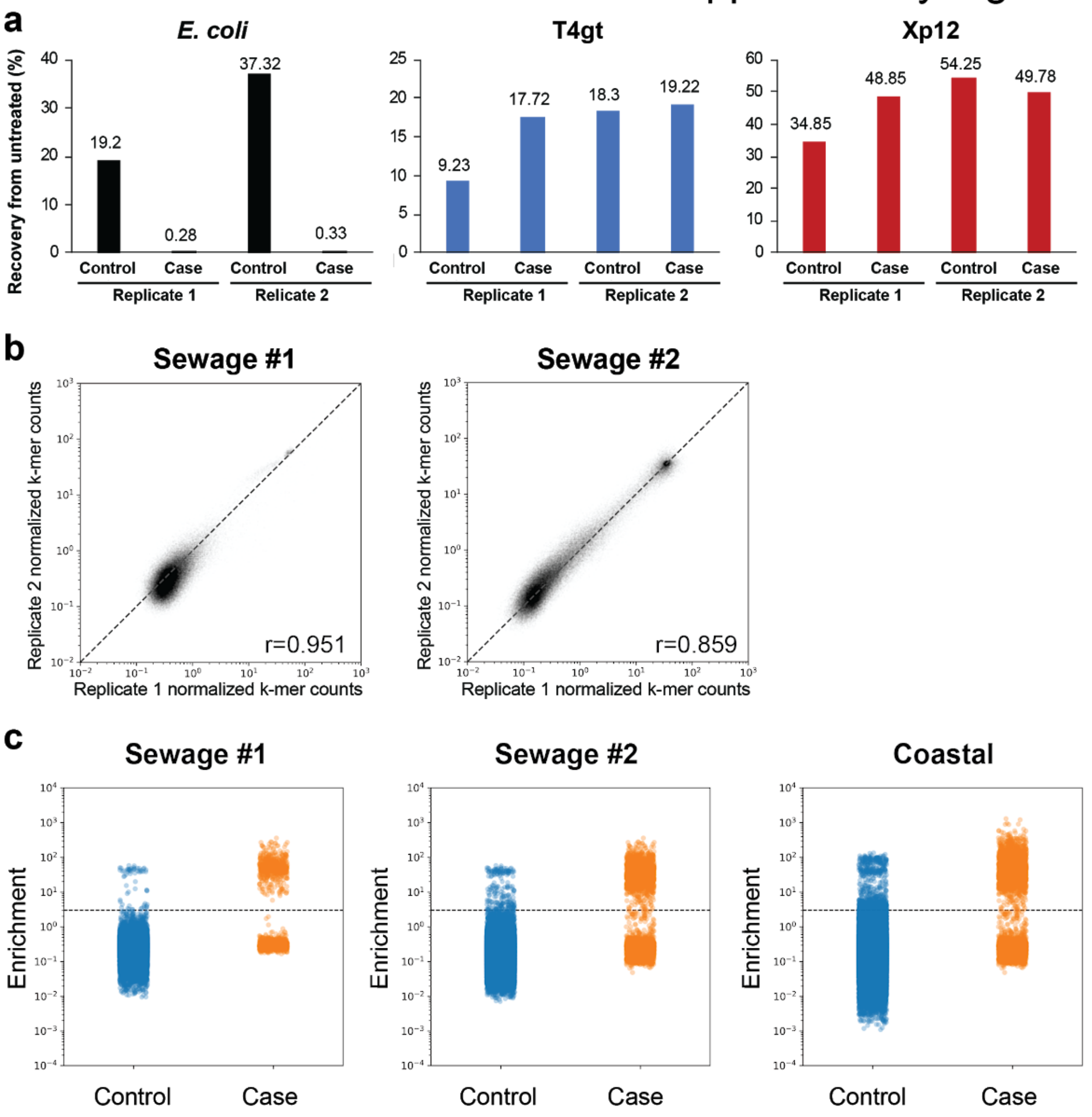


d

| Experiment | Modified contigs (>1000bp) | Unmodified contigs (>1000bp) |
|---|---|---|
| Sewage_1 | 660 | 72587 |
| Sewage_2 | 1905 | 81986 |
| Ocean | 1336 | 35802 |
| Total | 3901 | 190375 |

**Supplementary Figure 2** Validation of the selection method for DNA cytosine modification. **a**, Recoveries of various spiked-in DNA measured using qPCR quantification. Genomic DNA from *E. coli* (dC), XP12 (5mdC) and T4gt (5hmdC) were mixed at 1:1:1 ratio by molarity and a total of 100 ng DNA were spiked in the sequencing libraries from sewage #2. Recovery percentages of control (USER) and selection (TET2/BGT+APOBEC3A+USER) were normalized to untreated DNA. **b**, Correlation between two replicates from sewage samples #1 and #2. Each dot represents a unique k-mer. Scatter plots show normalized k-mer counts in each replicate. Pearson correlation coefficients (r) were calculated for each comparison. **c,** Strip plot showing the distribution of enrichment scores for each non-redundant contig larger than 1kb in sewage #1 (left) sewage #2 (center) and coastal samples (right). Dots represent the enrichment scores of individual non-redundant contigs. Enrichment score >=3 defines modified contigs while enrichment score < 3 defines unmodified contigs. **d**, Summary tables showing the number of modified and unmodified non-redundant contigs larger than 1 kb in each sample.

### Supplementary Figure 3

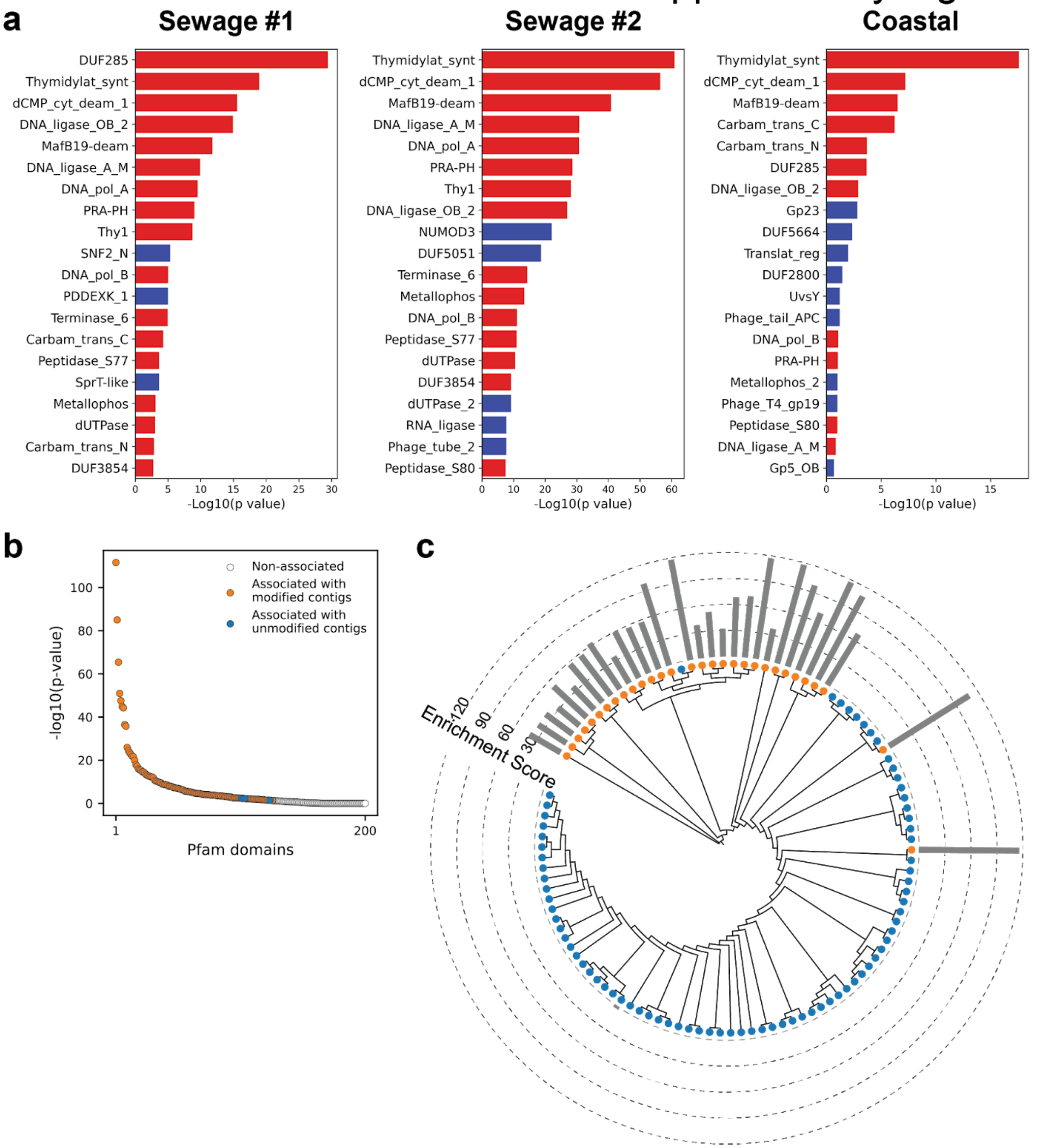

### Supplementary Figure 3

**d**

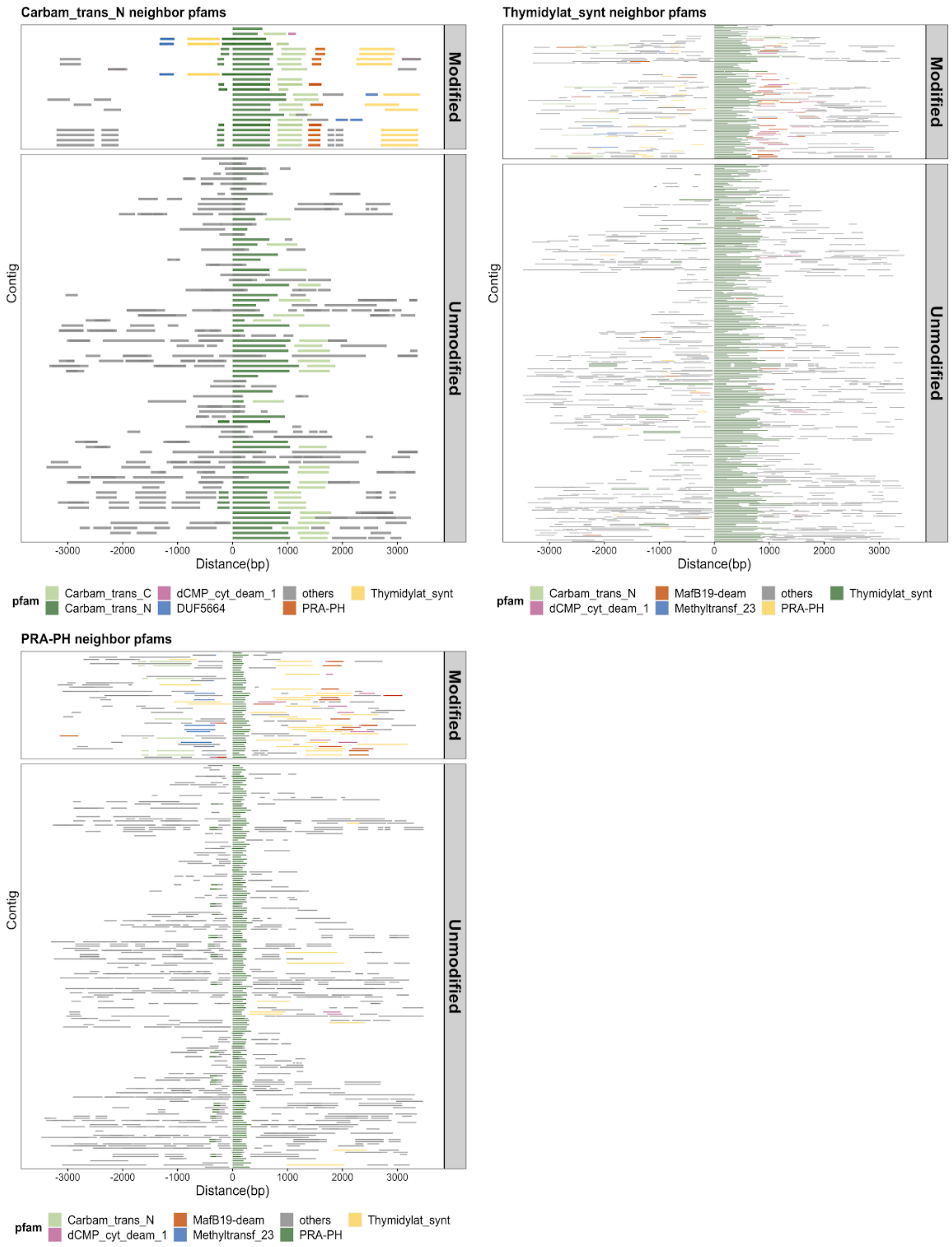

**Supplementary Figure 3** MetaGPA study reveals significantly associated Pfam domain. **a**, Top Pfam domains significantly associated with modified contigs (Bonferroni-corrected *p*-value <0.01) in each environment samples (two sewage and one coastal communities). Domains found significantly associated with modified contigs in at least two experiments were marked in red, blue denotes Pfam domains found significantly associated with modified contigs only in a single experiment. **b**, P value ranking of all Pfam domains in the composite dataset. **c**, Phylogenetic tree of carbamoyltransferase N-terminus. Orange and blue dots represent modified and unmodified contigs accordingly. Grey bars in the outer ring represent the enrichment scores of each contig with dashed lines showing the scales. Most of the carbamoyltransferase N-terminus domains from modified contigs form a distinctive phylogenetic branch. **d**, Genomic contexts of carbamoyltransferase N-terminus (top left panel), thymidylate synthase (top right panel) and phosphoribosyl-ATP pyrophosphohydrolase (PRA-PH, bottom panel) reveals commonly co-occurring Pfam domains in modified contigs. Neighboring Pfam domains spanning 3 kb upstream and downstream of the Pfam domain are displayed as solid lines. Top five co-occurring domains in modified contigs are colored and labelled. These co-occurring domains in modified contigs are, for the most part, absent in the unmodified contigs.

Supplementary Figure 4

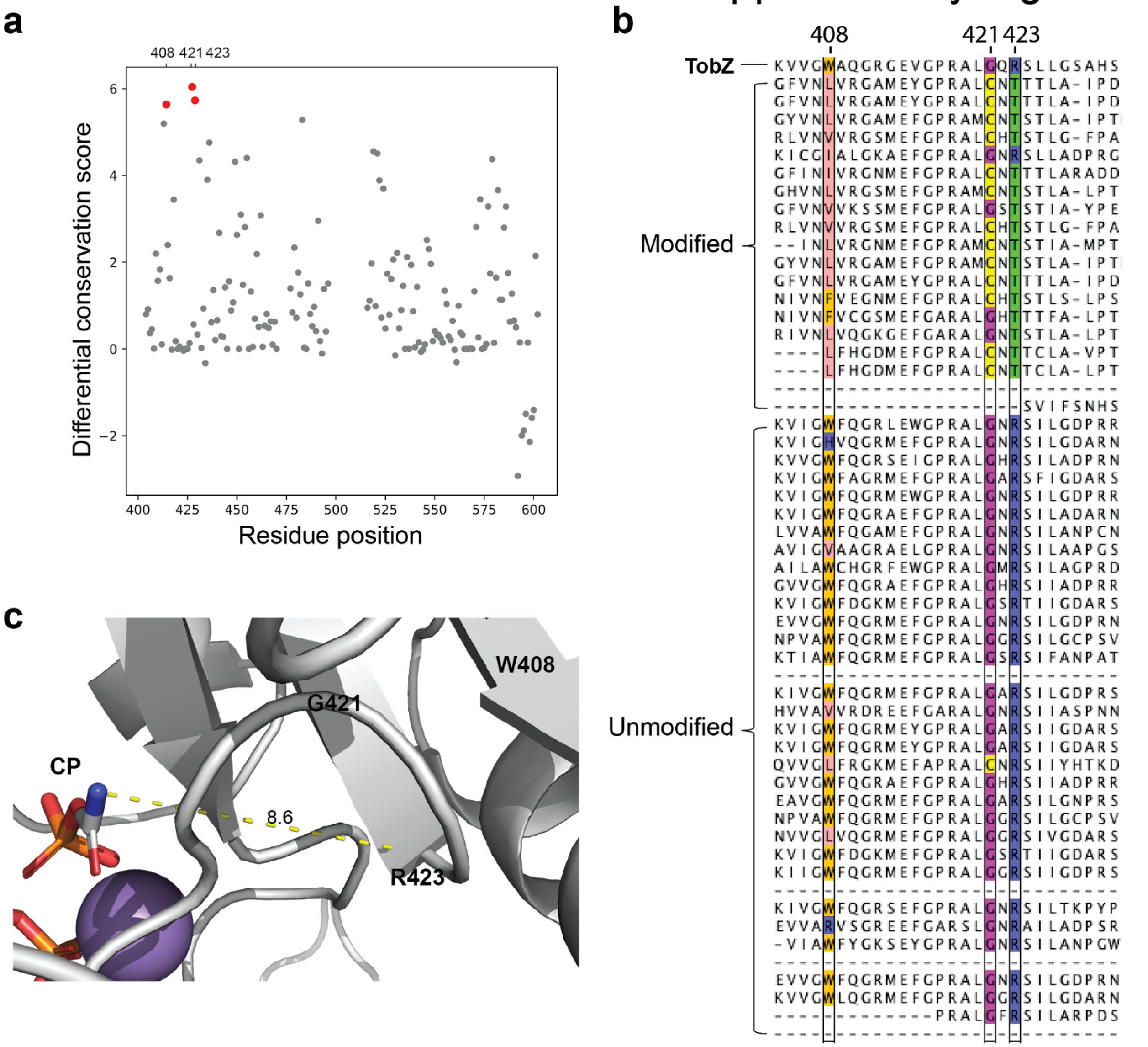


**Supplementary Figure 5** Association between key residues in the carbamoyltransferase C-terminus domains and DNA modification. **a**, Differential conservation score plotted against residue positions of the O-carbamoyltransferase TobZ. **b**, Multiple sequence alignment of 19 and 35 carbamoyltransferases from the modified and unmodified contigs respectively together with Tobz. Only part of the carbamoyltransferase C-terminus domain containing key residues are shown. TobZ Residues W408, G421 and R423 have been substituted for the most part with L, C and T in carbamoyltransferases found in modified contigs. These key residues are colored according to the type of amino acid. **c**, Structure of TobZ (PDB 3VET) highlighting M408, G421 and R423. Distance between R423 and carbamoyl phosphate (CP) is indicated in the picture (unit = Å).

### Supplementary Figure 5

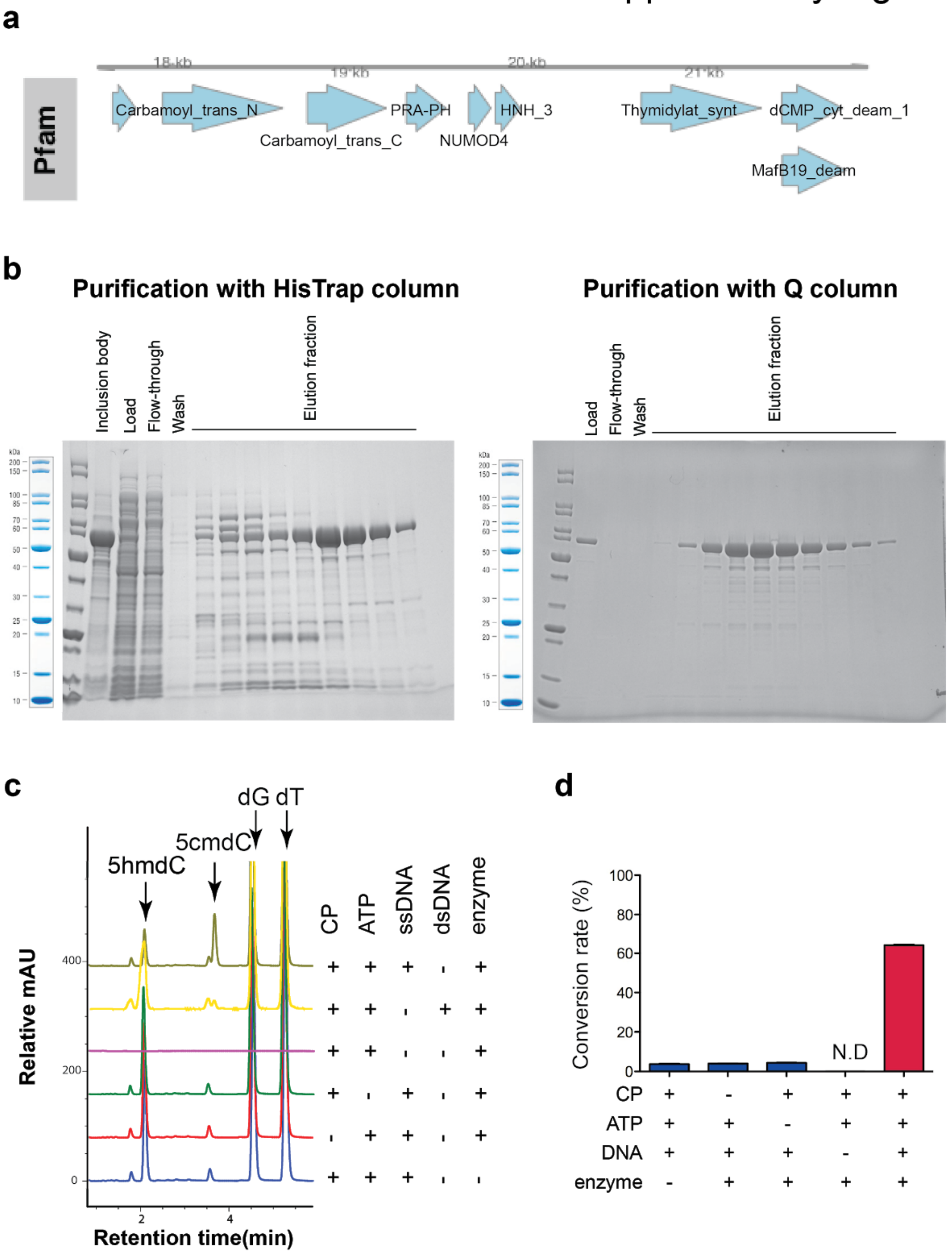

### Supplementary Figure 5

**e**

#### 5hmdC

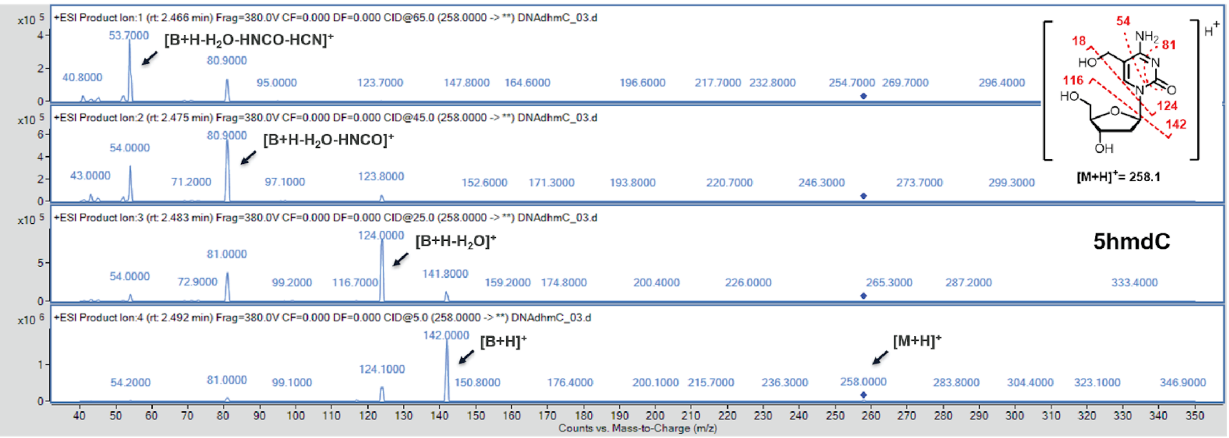


#### 5cmdC

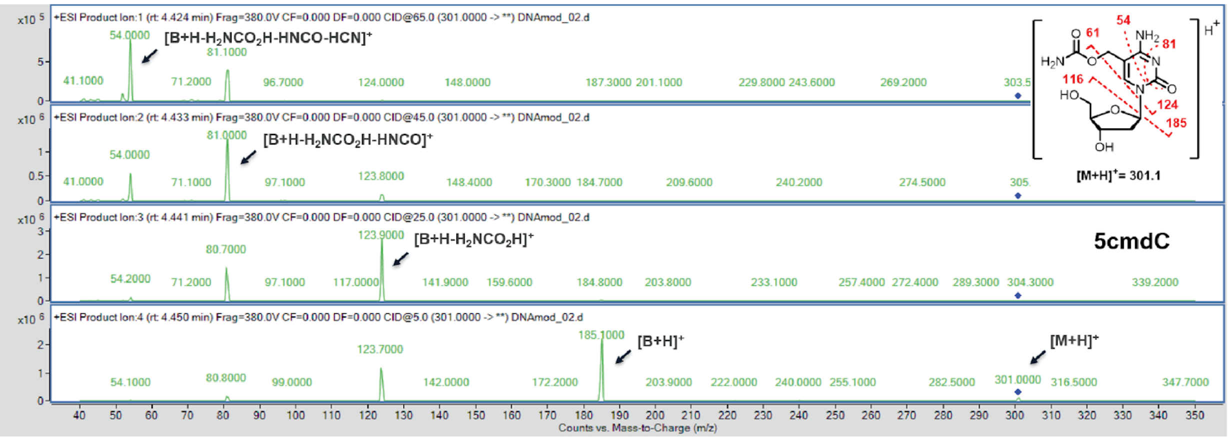

### Supplementary Figure 5

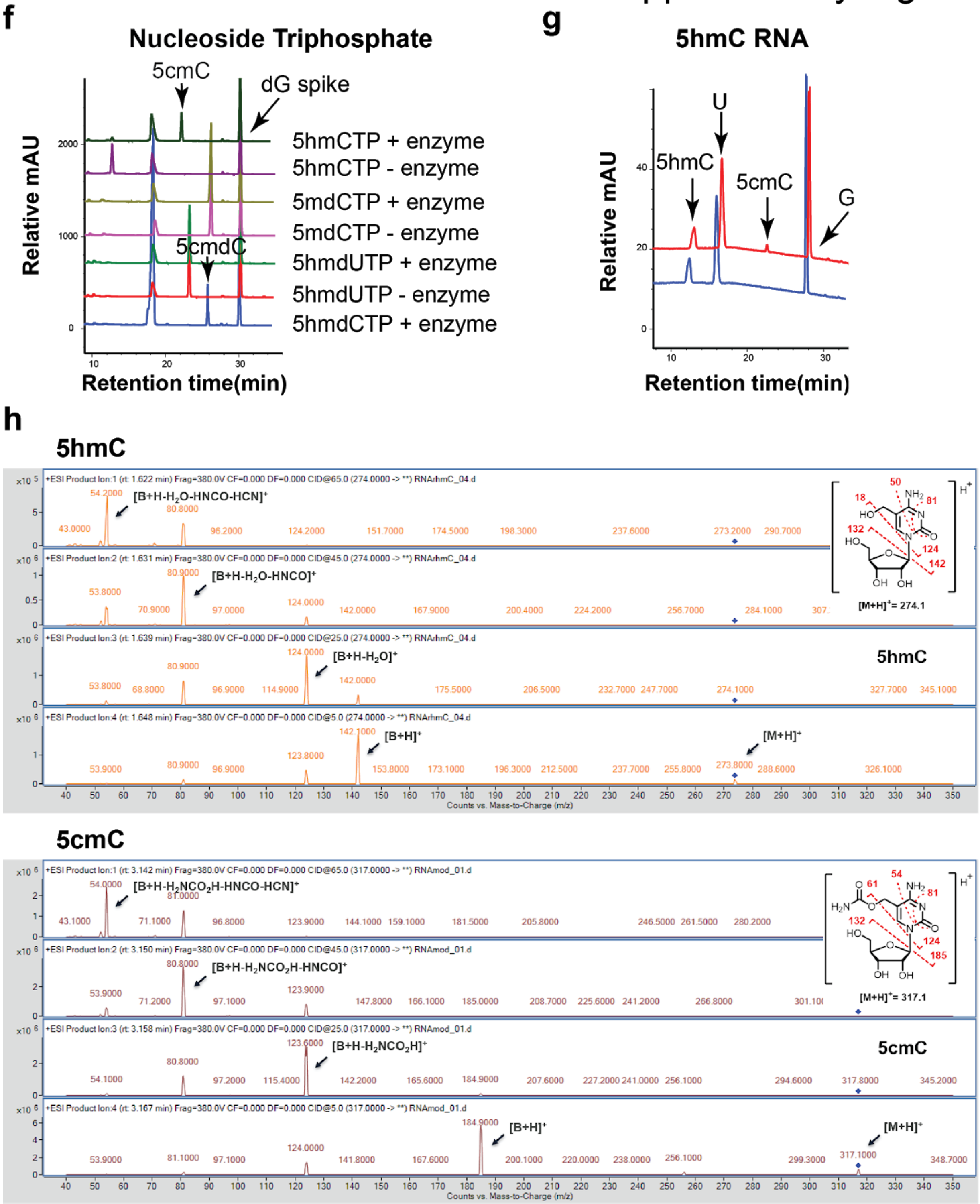


**Supplementary Figure 6** Validation of the novel 5-hydroxymethylcytosine carbamoyltransferase. **a**, Native genomic organization of the cloned carbamoyltransferase. **b**, SDS-PAGE gel showing purification of the carbamoyltransferase after two columns purification. Left, after HisTrap column purification ; right, after HisTrap and Q column purification. **c**, Characterization of enzymatic reaction by LC-MS. The arrow indicates the product peak 5cmdC. **d**, Quantification of enzymatic reactions from **c**. T4gt genomic DNA was used as a source of 5hmdC DNA. Bar graph represents average conversion rates ± SEM from three independent experiments. (N.D., not detected). **e**, + ESI-MS/MS fragmentation spectra of 5hmdC (top panel) and 5cmdC (bottom panel). **f**, Substrate selectivity. Enzymatic assays were performed with the following nucleoside triphosphates: 5hmdCTP, 5hmdUTP, 5mdCTP, and 5hmCTP. Carbamoylation was observed with 5hmdCTP and 5hmCTP (arrows). **g**, LC-MS trace for enzymatic reaction with a single-stranded RNA containing an internal 5hmC. **h**, + ESI-MS/MS fragmentation spectra of 5hmC (top panel) and 5cmC (bottom panel). Oligonucleotides and nucleoside triphosphate were converted to nucleosides before LC-MS analysis. A dG spike was added as internal reference for quantification of nucleoside triphosphates.

| Molarity Ratio | Modified DNA (ng) | Total DNA (ng) |
|---:|---:|---:|
| 1:3 | 8.639 | 250 |
| 1:10 | 1.127 | 250 |
| 1:100 | 0.093 | 250 |
| 1:1000 | 0.009 | 250 |

**Supplementary Table 1** Amounts of DNA used for enzymatic selection sensitivity test

**Supplementary Table 2** Pfam associations with DNA modification
